# The decline of global pollinator biodiversity in the Anthropocene

**DOI:** 10.1101/2024.12.22.629143

**Authors:** Ratheesh Kallivalappil, Sheena C. Cotter

## Abstract

Pollination underlies the functioning of ecosystems globally. Therefore, the endangerment and extinction of pollinator species are predicted to trigger cascade effects with the potential to alter the demographic collapse of complex ecological networks. However, although some studies have investigated the endangerment levels of pollinator species, the lack of global-scale analyses providing a universal overview of their extinction risks remains a major pending challenge in a world where climate change is rapidly decimating biodiversity. Here, we present the first truly global study of the endangerment level of vertebrate pollinators from across the tree of life. Based on a 1,666 species dataset, we investigate the macroecological patterns of species diversity and extinction risk of bird, mammal, and reptile pollinators of the world. We found higher extinction risk for mammal relative to bird and reptile pollinators. Globally, 1 in 3 mammal pollinators are currently threatened with extinction than 1 in 12 bird and 1 in 8 reptile pollinators. The hotspots of threatened bird pollinators mostly confined to Colombia and Hawaiian Islands, whereas the hotspots of threatened mammal and reptile pollinators are confined to Madagascar and various isolated islands. Notably, the endemic pollinators are more threatened than the widespread pollinators. The increasing decline of population will alter the status of threatened pollinators in future. While highlighting the quantity of threat and decline, we also show that the evolutionary predisposition along with habitat destruction for agriculture, and exploitation for bushmeat and pet trade are combinedly eroding the vertebrate pollinators biodiversity across the world at multiple scales. We suggest special environmental priorities such as controlled land-use, legislation on hunting, collaborative efforts between various stakeholders and community outreach programmes are essential for effective conservation of vertebrate pollinators.

## Introduction

The biosphere is entering a period of greatly increased extinction of species (Tittensor et al. 2014) with extinction rates several hundred times higher than the background rate (Barnosky et al. 2011). This growing rate of extinction calls for an urgent need for species’ conservation, as loss of species from an ecosystem leads to loss of the functional roles they play (Estes et al. 2011). A growing human population and associated environmental changes are increasing the risk of extinction for species around the world (Butchart et al. 2010, Hoffmann et al. 2010, Ripple et al. 2017). Therefore, identifying the patterns and drivers of biodiversity loss and extinction risk in species is crucial for conservation biology (Didham et al. 2007, Duenas et al. 2021).

Investigating extinction risk of pollinators is particularly important in the contemporary scenario, as pollination is one of the fundamental forms of ecological interactions which underlying the stability of ecosystem functioning on Earth (Kremen et al. 2007, Ollerton 2017, van der Plas 2019, Lemanski et al. 2022). Approximately 85% of angiosperm plants are pollinated by animals around the world (Ollerton et al. 2011), and pollinators are needed to pollinate 70% of the global food crops which humans consume (Klein et al. 2007). Therefore, the loss of pollinators can impact, other than just plant diversity, the associated organisms and a significant fraction of global agriculture, hence, human food security (Ollerton 2017, Lemanski et al. 2022).

Vertebrate pollination is widely recognized across the tropics (Fleming and Muchhala 2008, Fleming et al. 2009, Ratto et al. 2018). Therefore, the dependence of plants on vertebrate pollinators is higher in the tropics than at temperate or higher latitudes (Ford et al. 1979, Ford 1985, Traveset and Sáez 1997, Funamoto and Sugiura 2017, Ratto et al. 2018). They are necessary to maintain the tropical forests across the world, especially on islands (Marshall 1985, Cox et al. 1991, Elmqvist et al. 1992). For example, hummingbirds have a complex role in the diversification of plant species across tropical America, where they pollinate around 7,000 plant species of varying growth forms from ∼100 plant families (Leimberger et al. 2022, Barreto et al. 2024). In lowland tropical rain forest at La Selva, Costa Rica, it is estimated that around 25% of understorey plant species are pollinated by vertebrate species (Kress and Beach 1994). Similarly, a study in Malaysian old-growth forest revealed that 13.7% of trees were at least partially pollinated by vertebrates (Hodgkison et al. 2003). This was typically 3 – 11% per habitat at a mid-elevation tropical wet forest site at Western Ghats, India (Devy and Davidar 2003). Among vertebrates, birds are known to pollinate 52 of the 960 cultivated plants (Nabhan and Buchmann 1997), and typically they pollinate 5% of a region’s flora or 10% of flora if that region is an island (Anderson 2003, Kato and Kawakita 2004, Bernardello et al. 2006). Mammals, particularly bats are probably the main pollinators of approximately 1000 plants species across the tropics including several economically important crops such as agave, durian, dragon fruits, beans, and pitayas (Bumrungsri et al. 2008, Bumrungsri, 2009, Lobova et al. 2009, Kunz et al. 2011, Hernández and Salazar 2012, Aziz et al. 2017, Sritongchuay et al. 2019, Tremlett et al. 2020). The scattered studies on the economic value of vertebrate pollination services indicate that their important role as pollinators of tropical crops and the economy (Fujita and Tuttle 1991, Trejo-Salazar et al. 2016, Sheherazade et al. 2019). Pollination by reptiles is commonly recognized as an island phenomenon (Olesen and Valido 2003), where they mostly act as double mutualist: i.e., pollinate and disperse seeds of the same plant (Hansen and Müller 2009). This a risky prerequisite to the survival of island biota because they could often put the ecosystem in a vulnerable state when an interacting species extinct (Olesen et al. 2018, Fuster and Traveset 2019, Kahnt et al. 2023).

Despite the importance of pollinators for natural ecosystems and food production, several growing evidence points to substantial losses of pollinators in many regions across the globe (Buchmann and Nabhan 1996, Lennartsson 2002, Biesmeijer et al. 2006, Potts et al. 2010, Cameron et al. 2011, Dupont et al. 2011, Goulson et al. 2015, Powney et al. 2019, Vasiliev and Greenwood 2021). However, an increasing degree of attention is being given to the insect pollinators relative to vertebrate pollinators regardless of their crucial role in wild and crop plants pollination (Fujita and Tuttle 1991, Itino et al. 1991, Kunz et al. 2011, Ollerton 2017, Stewart and Dudash 2017, Ratto et al. 2018, Tremlett et al. 2020, Newman et al. 2021). In contrast to this, Regan et al. 2015 reported an early warning of bird and mammal pollinators biodiversity deterioration. They show each year, on average, 2.5 species of vertebrate pollinators moving one Red List category closer to extinction. However, this assessment was taxonomically and methodologically limited to bird and mammal pollinators using the Red List Index. But they did not quantitatively estimate the patterns of extinction risk and population decline and associated geographic and phylogenetic variations in extinction risk. Hence, empirical research needs to close some significant major gaps in knowledge about the pattern of current vertebrate pollinator diversity, spatial distribution, extinction hotspots, phylogenetic clustering in threat (i.e., the tendency of related groups to contain threatened taxa), and anthropogenic impact on extinction risk, especially at large taxonomic and geographic scales. To explore the big picture of pollinator distribution and extinction risk across geography, we ask the following questions: (i) Are there any geographical patterns in pollinators distributions and extinction risk? (ii) what proportion of bird, mammal and reptile pollinators are currently threatened across the globe? (iii) how does the pattern of pollinators’ threat and decline differ across the taxa? (iv) is there any nonrandom pattern in vertebrate pollinators extinction risk? and (v) how do anthropogenic factors threaten the pollinators’ biodiversity, and are there any patterns in their spatial distributions across the world?

## Materials and methods

### Pollinator species data collection

To extract pollinator species, we searched for published studies in ISI Web of Science and Google Scholar. We also searched the reference lists of all relevant papers, Biodiversity Heritage Library (biodiversitylibrary.org) and emailed the authors where papers could not be accessed for additional sources. We followed the criteria adopted by Regan et al. 2015 for the eligibility of species to be considered as a pollinator (see Supplementary Table S1). We searched for studies including a combination of the terms, for example, vertebrate polli*, bird polli*, avian polli*, passerine bird polli*, mammal polli*, bat polli*, reptile polli*, lizard polli*, ornithoph*, vertebrate flowe*, bird flowe*, mammal flowe*, bat flowe*, reptile flowe*, lizard flowe*, bird plant reproduct*, bat plant reproduct*, vertebrate plant interact*, bird plant interact*, mammal plant interact*, bat plant interact*, reptile plant interact*, lizard plant interact*, vertebrate flowe* reprod*, bird flowe* reprod*, mammal flowe* reprod*, bat flowe* reprod* in Web of Science; and vertebrate pollination, bird pollination, avian pollination, passerine bird pollination, mammal pollination, bat pollination, lizard pollination, reptile pollination, respectively in Google Scholar to narrow the search to pertinent studies.

Our exploration on the Web of Science yielded 21,677 papers; nevertheless, by narrowing our search to the initial 30 pages of Google Scholar, we obtained 2,400 articles (see similar method Kallivalappil et al. 2024) (see Table S2). Thus, a total of 24,077 papers were acquired from databases, with an additional 10 articles from alternate sources (Biodiversity Heritage Library and authors). Each paper’s title and abstract underwent meticulous scrutiny for significance, resulting in the evaluation of 4,147 studies for eligibility. Subsequently, after eliminating duplicative studies and thoroughly examining the full text of remaining articles, we identified 322 papers deemed suitable for our study. Utilizing these studies, we compiled our vertebrate pollinator database (see Figure S1 for the comprehensive PRISMA report). Our search focused on articles published until December 31, 2022. It’s important to note that our review was confined to English publications, thus introducing a bias towards non-English publications.

In our study we exclusively focused on extant species, thus meticulously refining our global dataset by excluding extinct and taxonomically unrecognized species. For example, we eliminated 17 bird and 1 mammal species (see Table S3) as extinct from the dataset of Regan et al. 2015. Though we noticed and corrected the taxonomic changes for 7 bird (*Celeus grammicus* to *Celeus undatus*, *Thraupis bonariensis* to *Pipraeidea bonariensis*; and *Zosterops montanus* combined with *Zosterops japonicus, Lichmera limbata* combined with *Lichmera indistincta*, *Philemon novaeguineae* combined with *Philemon buceroides* and *Arachnothera everetti* and *Heliangelus zusii* are no longer considered a species) and 3 mammal (*Artibeus incomitatus* combined with *Dermanura watsoni*; *Pteropus yapensis* and *Trichosurus arnhemensis* became subspecies of *Pteropus pelewensis* and *Trichosurus vulpecula*) pollinator species prior to our analyses. Additionally, we identified duplicate values for 3 mammal pollinators (*Dobsonia moluccensis*, *Mirza coquereli*, *Parantechinus apicalis*) of the 343 mammal pollinator species recorded in Regan et al. 2015 study, hence, we removed them from our dataset. We also omitted the species *Pycnonotus tricolor* (Diller et al. 2019) from our dataset due to unrecognised status of taxonomy by the BirdLife International. Thus, after cleaning the species, the dataset yielded an extant 1,255 bird, 371 mammal and 40 reptile pollinator species for the analyses (see Supplementary Material: Pollinator Dataset). We followed the taxonomy of the Handbook of the Birds of the World and BirdLife International (BirdLife International 2023) for bird pollinators; the IUCN Red List mammal taxonomy (IUCN 2023) for mammal pollinators; and the reptile database (http://www.reptile-database.org; Uetz et al. 2023) for reptile pollinators as the taxonomic authority.

### Assessing extinction risk and population trends of the world’s pollinators

We retrieved the IUCN risk status and population trend data for our species using *rl_search* from the package ***rredlist*** (Gearty and Chamberlain 2022) to determine variation in the degrees of extinction risks and population decline of the world’s pollinators across the described taxon. To quantify variation in extinction risk across groups with each of these categories, we employed the consensus approach (Clausnitzer et al. 2009, Hoffmann et al. 2010, Böhm et al. 2013, Pincheira-Donoso et al. 2021, Pincheira-Donoso et al. 2023, Kallivalappil et al. 2024) which calculate proportions of threatened (Critically Endangered, Endangered and Vulnerable) species, while also accounting for the uncertainty of Data Deficient species, which were assumed to fall into the above threatened categories in the same proportions as non-Data Deficient species, according to the formula: Prop_threat_ = (CR + EN + VU)/(*N* – DD), where *N* is the total number of species in the sample per category, CR, EN and VU are the numbers of species in each of the above IUCN categories, and DD is the number of species in the Data Deficient category. However, there is uncertainty associated with the assumption that DD species will fall across threatened categories in similar proportions and thus, future assessments of these species may result in an alteration of these estimates (Böhm et al. 2013). Therefore, to counterbalance potential uncertainties, we calculated the upper and lower bounds of the degrees of threatened species by assuming that (*i)* no DD species fall into one of the threatened categories [lower margin; Prop_threat_ = (CR + EN + VU)/(N)], and that (*ii*) all DD species fall into a threatened category [upper margin; Prop _threat_ = (CR + EN + VU + DD)/(N)] (Böhm et al. 2013).

We further estimated the proportion of species with decreasing populations as: PropDecr = DecreaseN/(N − UnknownN) where DecreaseN is the number of decreasing species, UnknownN is the number of unknown-trend species, and N is the total number of species in all four population trends (Finn et al. 2023, Kallivalappil et al. 2024). We accounted for species with unknown population trends in our estimate, the same as DD species accounted for in the estimation of proportion of threat (Finn et al. 2023, Kallivalappil et al. 2024). We estimated the upper and lower bound of population trend by assuming that no unknown-trend species are decreasing [lower margin; PropDecrL = DecreaseN/N] but assumes all unknown-trend species are decreasing [upper margin; PropDecrU = (DecreaseN + UnknownN)/N]. We estimated the current, upper, and lower limits for species with stable and increasing population trends also. We estimated this for both pollinators (birds, mammals, reptiles) and global species (birds, mammal, reptiles), and separately for the major families of bird and mammal pollinator groups (i.e., for birds, the categorization was based on the family if contains a minimum of 27 or more species, but for mammals, if the family contains a minimum of 100 species).

### Spatial patterns of species richness

To explore the spatial distribution of pollinator richness, we obtained all available spatial distribution data (in the form of shapefiles) for mammal pollinators from the IUCN Red List (International Union for Conservation of Nature; https://www.iucnredlist.org; downloaded on 25/07/23) advanced search criteria portal. However, we obtained spatial data of reptile pollinators by selecting specific taxonomic groups from the same portal (downloaded on 28/09/23). For birds, we obtained all available spatial data from the BirdLife International portal on request (BirdLife International 2023; downloaded on 28/07/23). We imported these spatial data into R software (version 4.2.3; R Core Team 2023) using the function ***read_sf*** from the **sf** package (Pebesma 2018) and then extracted the pollinator species’ shapefiles using the function ***filter*** from the same package. As a result, we obtained complete spatial data for mammal pollinators (total 371), but we obtained spatial data for 1,253 bird (total of 1255) and 39 reptile (total of 40) pollinators only.

We then cleaned each species’ shapefile by using the function ***lets.shFilter*** from the package **letsR** (Vilela and Villalobos 2015). We considered the polygons of species spatial data with the following IUCN map code criteria indicated as **Presence** = *extant* (code 1); **Origin** = *native* (code 1) and *reintroduced* (code 2); and **Seasonal** = *resident* (code 1), *breeding* (code 2) and *non-breeding* (code 3) [Presence (*extant*) + Origin (*native and reintroduced*) + Seasonal (*resident, breeding and non-breeding*)] (see similar method Joppa et al. 2016). After cleaning, we retrieved polygons data for 1,249 birds (total of 1,253 species), 369 mammals (total of 371 species) and 39 reptiles (total of 40 species) pollinator species.

However, the data in this format was not available for some of our pollinators. Therefore, for four bird pollinators we used the polygons coded as *possibly extant* (code 3) of **Presence** for *Discosura letitiae*, and *possibly extinct* (code 4) of **Presence** for *Charmosyna diadema*, *Eriocnemis godini*, and *Psittirostra psittacea* with *native* (code 1) of **Origin** and *resident* (code 1) of **Seasonal** codes: [Presence (*possibly extant*) + Origin (*native*) + Seasonal (*resident*)]; [Presence (*possibly extinct*) + Origin (*native*) + Seasonal (*resident*)].

For two mammal pollinators, we used the polygons coded as *possibly extinct* (code 4; *Mystacina robusta*) and *presence uncertain* (code 6; *Pteropus melanotus*) of **Presence** along with *native* (code 1) of **Origin** and *resident* (code 1) of **Seasonal**: [Presence (*possibly extinct*) + Origin (*native*) + Seasonal (*resident*)]; [Presence (*presence uncertain*) + Origin (*native*) + Seasonal (*resident*)].

For reptile pollinator *Cnemidophorus murinus*, we used the polygon coded as *origin uncertain* (code 5) of **Origin** with *extant* (code 1) of **Presence** and *resident* (code 1) of **Seasonal** codes: [Presence (*extant*) + Origin (*origin uncertain*) + Seasonal (*resident*)].

Moreover, the spatial data for two bird (*Chloropsis cochinchinensis*, *Zosterops flavus*) and a reptile pollinator species (*Pseudocordylus subviridis*) were unavailable in the IUCN portal, therefore, we obtained them from the Map of Life portal (https://mol.org/species/). We also obtained spatial data for the reptile pollinator species, *Cyclura carinata*, from the same portal because we omitted the IUCN spatial data for this species due to the issue of overestimation of species richness. We spatially joined these data with the IUCN and BirdLife International spatial data by matching the field attributes using the software R. Hence, we obtained complete spatial data for all pollinator taxa for further analysis.

To explore the spatial patterns of species richness distributions, we created species richness maps by spatially joining the spatial data with 0.5° by 0.5° latitude-longitude gridded (approximately 50 km at the Equator) global map using the function ***st_join*** from the **sf** package. We counted the number of overlapping polygons for each grid cell, resulting in a species richness count (see method, Kallivalappil et al. 2024).

To produce the richness maps, the bird and mammal pollinators were grouped into a wide range of categories according to the IUCN Red List criteria:

i. all species (birds = 1255, mammals = 371), species classified as threatened (birds = 104, mammals = 102), not-threatened (birds = 1151, mammals = 269), and data deficient (birds = 2, mammals = 16).
ii. Species whose population trend is decreasing (birds = 507, mammals = 152), stable (birds = 528, mammals = 117), increasing (birds = 71, mammals = 7), and unknown (birds = 149, mammals = 95), according to the IUCN.
iii. Species which threatened from anthropogenic impacts such as habitat destruction (birds = 194, mammals = 182), overexploitation (birds = 42, mammals = 119), climate change (birds = 134, mammals = 50), invasion (birds = 69, mammals = 31), food trade (birds = 93, mammals = 130), and pet trade (birds = 734, mammals = 25).
iv. Species with endemic status (species present only in a particular political boundary ; birds = 433, mammals = 101), and globally threatened birds (n = 1315) and mammals (n = 1318). But, for reptile pollinators, we restricted maps to (v) all species (n = 40), threatened (n = 5) and decreasing (n = 4) population trend.

We also produced maps of (vi) proportion of threatened bird and mammal pollinators. Using the function *match* from the R software, we transferred variables from the threatened shapefile to all species’ shapefile (0.5° gridded shapefile produced to map all species richness) and estimated the proportion for each grid cell by dividing the threatened species richness by all species’ richness. We visualized the richness patterns using the R package **tmap** (Tennekes 2018). All maps were created using the World Geodetic System (WGS84) coordinates.

### Assessing spatial variations in distribution of species richness

To examine the spatial variation between bird and mammal pollinators distribution, a Chi-square test in R (function *prop.test*) was used to compare the proportions of bird and mammal pollinators presence (minimum one species per grid cell) with the total number of grid cells (which we obtained from our species richness mapping).

### Patterns of spatial correlation between data deficient and threatened pollinators

To assess the potential threat posed to bird and mammal pollinators classified as data deficient in the IUCN database, we conducted an analysis of spatial association. This involved evaluating the relationship between the richness of threatened species and the richness of species with data deficient status, employing Tjøstheim’s coefficient with the function ***cor.spatial*** from the **SpatialPack** package (Vallejos et al. 2020). A positive correlation between them suggests the possibility of data deficient species facing similar anthropogenic threat across its range, same as threatened pollinators. Using the same function, we also evaluated the spatial association between the globally threatened bird and mammal species richness with that of pollinator bird and mammal richness.

### Anthropogenic factors of extinction risk

We retrieved the status of endemism, anthropogenic threats and use and trade data for bird, mammal and reptile pollinators manually from the IUCN portal.

### Phylogenetic patterns of extinction risk

To test the role of shared ancestry on species threat and population decline, we constructed maximum clade credibility tree (package **ape**; Paradis and Schliep 2019) using 1000 randomly selected phylogenies from the posterior distribution of phylogenies from Jetz et al. 2012 for bird and Upham et al. 2019 for mammal pollinators. We used the function ***phylo.signal.disc*** (Bascompte et al. 2019) to test the phylogenetic signals. We did not conduct this analysis for reptile pollinators due to low sample size and insufficient Red List status.

## Results

### Diversity of pollinators

Using the mainstream databases, we identified a diversity of 1,255 birds (12 order, 64 families), 371 mammals (10 order, 37 families) and 40 reptile species (1 order, 10 families) as vertebrate pollinators from the literature, a global representation of 11.4% bird (n = 11,024), 6.3% of mammal (n = 5,886) and 0.4% of reptile (n = 10,222) species (data retrieved from the IUCN portal). Our dataset for bird pollinators showed a diversity of 35% of bird pollinators (n = 433) as endemic to a particular political boundary. However, this was 27% for mammals (n = 101) and 78% for reptile (n = 31) pollinators.

### Extinction risk of world’s pollinators

Utilizing data from the IUCN, our evaluation of the extinction risk facing the world’s vertebrate pollinators revealed a pronounced asymmetry among various taxonomic groups, with certain groups exhibiting notably elevated levels of threat. Specifically, our findings indicated that mammal pollinators (29%) faced a significantly higher risk of extinction compared to bird pollinators (8.3%; see Table 1 & S5). Moreover, our analysis revealed that approximately 13% of reptile pollinators were threatened with extinction (see Table 1). Furthermore, we observed a significantly elevated threat level among endemic bird and mammal pollinators compared to their widespread counterparts (see Table 1 and Tables S6 & 7).

**Table 1.**
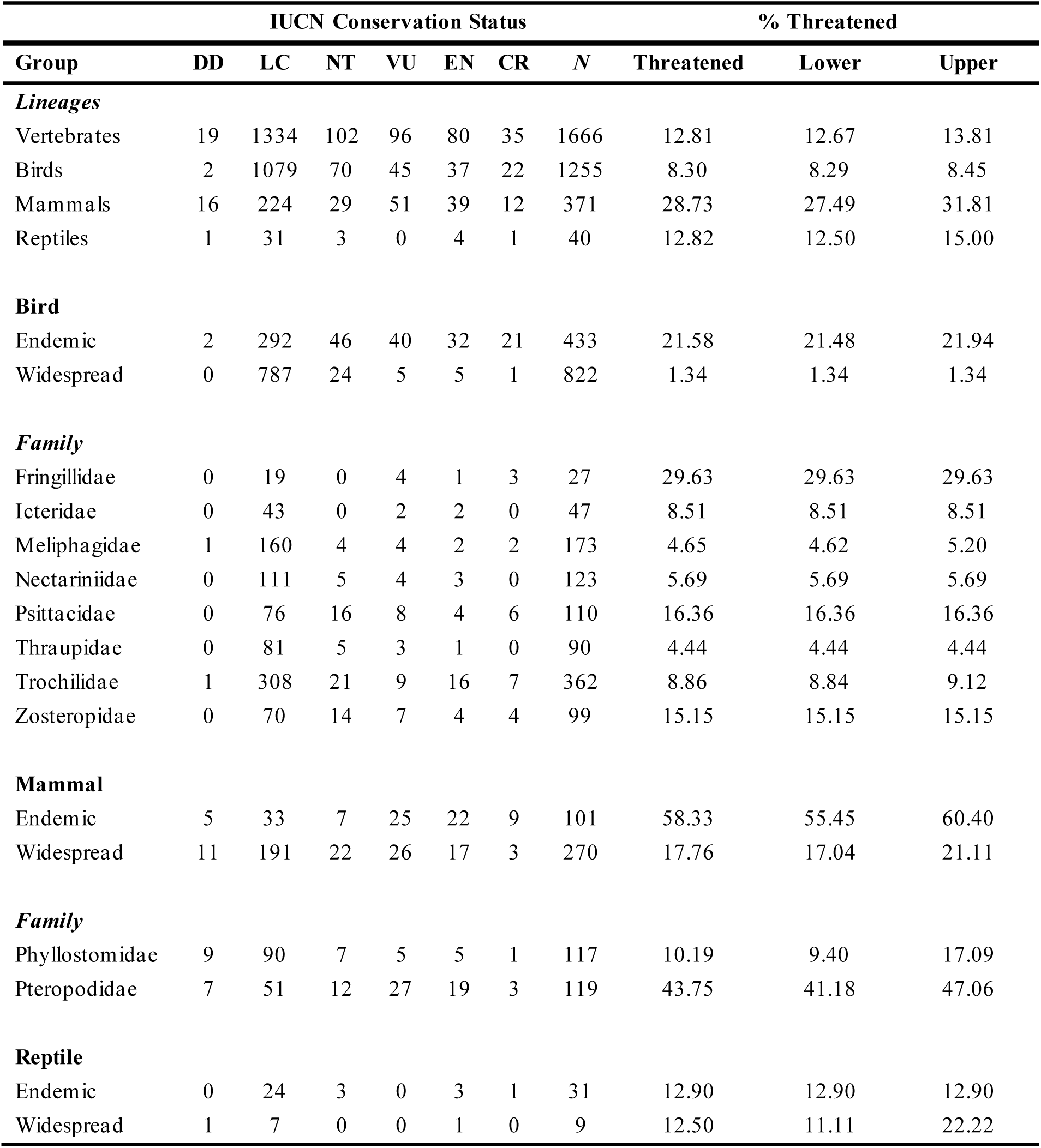
Extinction risk of the world’s vertebrate pollinators organized as major groups and broken down into major families within groups. Each IUCN Red List category indicates: DD = Data Deficient, LC = Least Concern, NT = Near Threatened, VU = Vulnerable, EN = Endangered, and CR = Critically Endangered. N is the total number of species in each group and the lower and upper level of threat are the proportion of species threatened by considering the Data Deficient species as not-threatened and threatened, according to global-scale measures produced by the IUCN.

| Group | IUCN Conservation Status |  |  |  |  |  |  | % Threatened |  |  |
| --- | --- | --- | --- | --- | --- | --- | --- | --- | --- | --- |
|  | DD | LC | NT | VU | EN | CR | N | Threatened | Lower | Upper |
| **Lineages** |  |  |  |  |  |  |  |  |  |  |
| Vertebrates | 19 | 1334 | 102 | 96 | 80 | 35 | 1666 | 12.81 | 12.67 | 13.81 |
| Birds | 2 | 1079 | 70 | 45 | 37 | 22 | 1255 | 8.30 | 8.29 | 8.45 |
| Mammals | 16 | 224 | 29 | 51 | 39 | 12 | 371 | 28.73 | 27.49 | 31.81 |
| Reptiles | 1 | 31 | 3 | 0 | 4 | 1 | 40 | 12.82 | 12.50 | 15.00 |
| **Bird** |  |  |  |  |  |  |  |  |  |  |
| Endemic | 2 | 292 | 46 | 40 | 32 | 21 | 433 | 21.58 | 21.48 | 21.94 |
| Widespread | 0 | 787 | 24 | 5 | 5 | 1 | 822 | 1.34 | 1.34 | 1.34 |
| **Family** |  |  |  |  |  |  |  |  |  |  |
| Fringillidae | 0 | 19 | 0 | 4 | 1 | 3 | 27 | 29.63 | 29.63 | 29.63 |
| Icteridae | 0 | 43 | 0 | 2 | 2 | 0 | 47 | 8.51 | 8.51 | 8.51 |
| Meliphagidae | 1 | 160 | 4 | 4 | 2 | 2 | 173 | 4.65 | 4.62 | 5.20 |
| Nectariniidae | 0 | 111 | 5 | 4 | 3 | 0 | 123 | 5.69 | 5.69 | 5.69 |
| Psittacidae | 0 | 76 | 16 | 8 | 4 | 6 | 110 | 16.36 | 16.36 | 16.36 |
| Thraupidae | 0 | 81 | 5 | 3 | 1 | 0 | 90 | 4.44 | 4.44 | 4.44 |
| Trochilidae | 1 | 308 | 21 | 9 | 16 | 7 | 362 | 8.86 | 8.84 | 9.12 |
| Zosteropidae | 0 | 70 | 14 | 7 | 4 | 4 | 99 | 15.15 | 15.15 | 15.15 |
| **Mammal** |  |  |  |  |  |  |  |  |  |  |
| Endemic | 5 | 33 | 7 | 25 | 22 | 9 | 101 | 58.33 | 55.45 | 60.40 |
| Widespread | 11 | 191 | 22 | 26 | 17 | 3 | 270 | 17.76 | 17.04 | 21.11 |
| **Family** |  |  |  |  |  |  |  |  |  |  |
| Phyllostomidae | 9 | 90 | 7 | 5 | 5 | 1 | 117 | 10.19 | 9.40 | 17.09 |
| Pteropodidae | 7 | 51 | 12 | 27 | 19 | 3 | 119 | 43.75 | 41.18 | 47.06 |
| **Reptile** |  |  |  |  |  |  |  |  |  |  |
| Endemic | 0 | 24 | 3 | 0 | 3 | 1 | 31 | 12.90 | 12.90 | 12.90 |
| Widespread | 1 | 7 | 0 | 0 | 1 | 0 | 9 | 12.50 | 11.11 | 22.22 |

Our analyses concerning the extinction risk facing bird families revealed a notable proportion of threat within the Fringillidae family (30%; see Table 1). Conversely, for mammals, a high level of threat was identified within the Pteropodidae family (44%; see Table 1). Furthermore, our findings indicated a correlation between family size (i.e., the number of species per family) and the number of threatened species within respective bird and mammal families (see Tables S8 & 9).

### Population trends of world’s pollinators

It is of significant concern that a substantial proportion of populations across all taxonomic groups are experiencing alarming declines. Our research revealed that half of the population of mammalian pollinators exhibited a significant decrease (55%), compared to bird pollinators - where the decrease was slightly lower at 46% (see Table 2 & S10). Notably, reptile pollinators experienced the least population decrease, with only 14% showing declines (see Table 2). Additionally, our analysis regarding population decreases among endemic and widespread pollinators revealed that a larger proportion of endemic bird (51%) and mammal (71%) pollinators are experiencing significant declines on a large-scale level (see Table 2 and S11 & 12). However, in reptiles, the population of widespread pollinators is declining more than the endemic pollinators (see Table 2).

**Table 2.** Patterns of population trends among the world’s vertebrate pollinators species organized as major groups and broken down into major families within groups. Each group shows the total number of species (N), and the proportion of species per group that are decreasing, stable, or increasing in population size along with their lower and upper levels of population trend by considering the Unknown trend species as not-threatened and threatened, according to global-scale measures produced by the IUCN.

| Group | IUCN Population Trends |  |  |  |  |  |  |  |  |  |  |  |  |  |
| --- | --- | --- | --- | --- | --- | --- | --- | --- | --- | --- | --- | --- | --- | --- |
|  | N | Unknown | Decreasing | Current | Lower | Upper | Stable | Current | Lower | Upper | Increasing | Current | Lower | Upper |
| Vertebrates | 1666 | 255 | 663 | 46.99 | 39.80 | 55.10 | 669 | 47.41 | 40.16 | 55.46 | 79 | 5.60 | 4.74 | 20.05 |
| Birds | 1255 | 149 | 507 | 45.84 | 40.40 | 52.27 | 528 | 47.74 | 42.07 | 53.94 | 71 | 6.42 | 5.66 | 17.53 |
| Mammals | 371 | 95 | 152 | 55.07 | 40.97 | 66.58 | 117 | 42.39 | 31.54 | 57.14 | 7 | 2.54 | 1.89 | 27.49 |
| Reptiles | 40 | 11 | 4 | 13.79 | 10.00 | 37.50 | 24 | 82.76 | 60.00 | 87.50 | 1 | 3.45 | 2.50 | 30.00 |
| **Bird** |  |  |  |  |  |  |  |  |  |  |  |  |  |  |
| Endemic | 433 | 57 | 193 | 51.33 | 44.57 | 57.74 | 163 | 43.35 | 37.64 | 50.81 | 20 | 5.32 | 4.62 | 17.78 |
| Widespread | 822 | 92 | 314 | 43.01 | 38.20 | 49.39 | 365 | 50.00 | 44.40 | 55.60 | 51 | 6.99 | 6.20 | 17.40 |
| *Family* |  |  |  |  |  |  |  |  |  |  |  |  |  |  |
| Fringillidae | 27 | 1 | 11 | 42.31 | 40.74 | 44.44 | 11 | 42.31 | 40.74 | 44.44 | 4 | 15.38 | 14.81 | 18.52 |
| Icteridae | 47 | 0 | 24 | 51.06 | 51.06 | 51.06 | 21 | 44.68 | 44.68 | 44.68 | 2 | 4.26 | 4.26 | 4.26 |
| Meliphagidae | 173 | 26 | 40 | 27.21 | 23.12 | 38.15 | 100 | 68.03 | 57.80 | 72.83 | 7 | 4.76 | 4.05 | 19.08 |
| Nectariniidae | 123 | 0 | 35 | 28.46 | 28.46 | 28.46 | 88 | 71.54 | 71.54 | 71.54 | 0 | 0.00 | 0.00 | 0.00 |
| Psittacidae | 110 | 1 | 56 | 51.38 | 50.91 | 51.82 | 45 | 41.28 | 40.91 | 41.82 | 8 | 7.34 | 7.27 | 8.18 |
| Thraupidae | 90 | 3 | 30 | 34.48 | 33.33 | 36.67 | 54 | 62.07 | 60.00 | 63.33 | 3 | 3.45 | 3.33 | 6.67 |
| Trochilidae | 362 | 69 | 187 | 63.82 | 51.66 | 70.72 | 97 | 33.11 | 26.80 | 45.86 | 9 | 3.07 | 2.49 | 21.55 |
| Zosteropidae | 99 | 29 | 50 | 71.43 | 50.51 | 79.80 | 18 | 25.71 | 18.18 | 47.47 | 2 | 2.86 | 2.02 | 31.31 |
| **Mammal** |  |  |  |  |  |  |  |  |  |  |  |  |  |  |
| Endemic | 101 | 17 | 60 | 71.43 | 59.41 | 76.24 | 22 | 26.19 | 21.78 | 38.61 | 2 | 2.38 | 1.98 | 18.81 |
| Widespread | 270 | 78 | 92 | 47.92 | 34.07 | 62.96 | 95 | 49.48 | 35.19 | 64.07 | 5 | 2.60 | 1.85 | 30.74 |
| *Family* |  |  |  |  |  |  |  |  |  |  |  |  |  |  |
| Phyllostomidae | 117 | 59 | 12 | 20.69 | 10.26 | 60.68 | 46 | 79.31 | 39.32 | 89.74 | 0 | 0.00 | 0.00 | 50.43 |
| Pteropodidae | 119 | 25 | 60 | 63.83 | 50.42 | 71.43 | 31 | 32.98 | 26.05 | 47.06 | 3 | 3.19 | 2.52 | 23.53 |
| **Reptile** |  |  |  |  |  |  |  |  |  |  |  |  |  |  |
| Endemic | 31 | 7 | 3 | 12.50 | 9.68 | 32.26 | 20 | 83.33 | 64.52 | 87.10 | 1 | 4.17 | 3.23 | 25.81 |
| Widespread | 9 | 4 | 1 | 20.00 | 11.11 | 55.56 | 4 | 80.00 | 44.44 | 88.89 | 0 | 0.00 | 0.00 | 44.44 |

Our findings revealed a diverse pattern of population decrease among bird and mammal pollinator families. Within bird families, a range of 27% to 71% of population decrease was identified, with more than half of the population of families such as Icteridae, Psittacidae, Trochilidae, and Zosteropidae experiencing declines worldwide (see Table 2). Conversely, in mammals, approximately 64% of the Pteropodidae population experienced a decrease (see Table 2). Similarly to the observed relationship between family size and the number of threatened species, our analysis also indicated a correlation between family size and the number of species experiencing population decreases within bird and mammal pollinator groups (see Tables S8 & 9).

### Taxonomic variations in threat and population trends of world’s pollinators

Our analyses to examine the relationship between the threat and population decrease at taxonomic level showed that mammalian pollinators were significantly more threatened than bird pollinators (*X^2^* = 101.57, *df* = 1, *p* = < 0.001). The same pattern was noticed with the population decrease where the population of mammalian pollinators decreased more (*X^2^* = 7.180, *df* = 1, *p* = 0.007). Analysis with the endemic and widespread pollinators showed that endemic bird and mammal pollinators were more threatened (Bird: *X^2^* = 149.53, *df* = 1, *p* = <0.001; Mammal: *X^2^* = 54.341, *df* = 1, *p* = <0.001) or decreasing (Bird: *X^2^* = 6.5821, *df* = 1, *p* = 0.010; Mammal: *X^2^* = 12.123, *df* = 1, *p* = 0.001) than to widespread pollinators.

### Spatial patterns of species richness

We observed an asymmetrical distributional pattern in the spatial richness of bird, mammal, and reptile pollinators. Specifically, bird pollinators exhibited a significantly broader distributional range compared to mammal pollinators (see Table S13). Notably, regions such as the tropical Andes and the Atlantic Forest emerged as prominent hotspots for both bird and mammal pollinators (see Figure 1a & b). Reptile pollinators, on the other hand, were predominantly restricted to the Socotra Island in Yemen and the northern part of New Zealand (see Figure S2a). Our analysis of endemic pollinator biodiversity richness revealed that the east or southeastern region of Australia harbours the largest hotspot of endemic bird and mammal pollinators’ richness (see Figure S3a & b).

**Figure 1.**
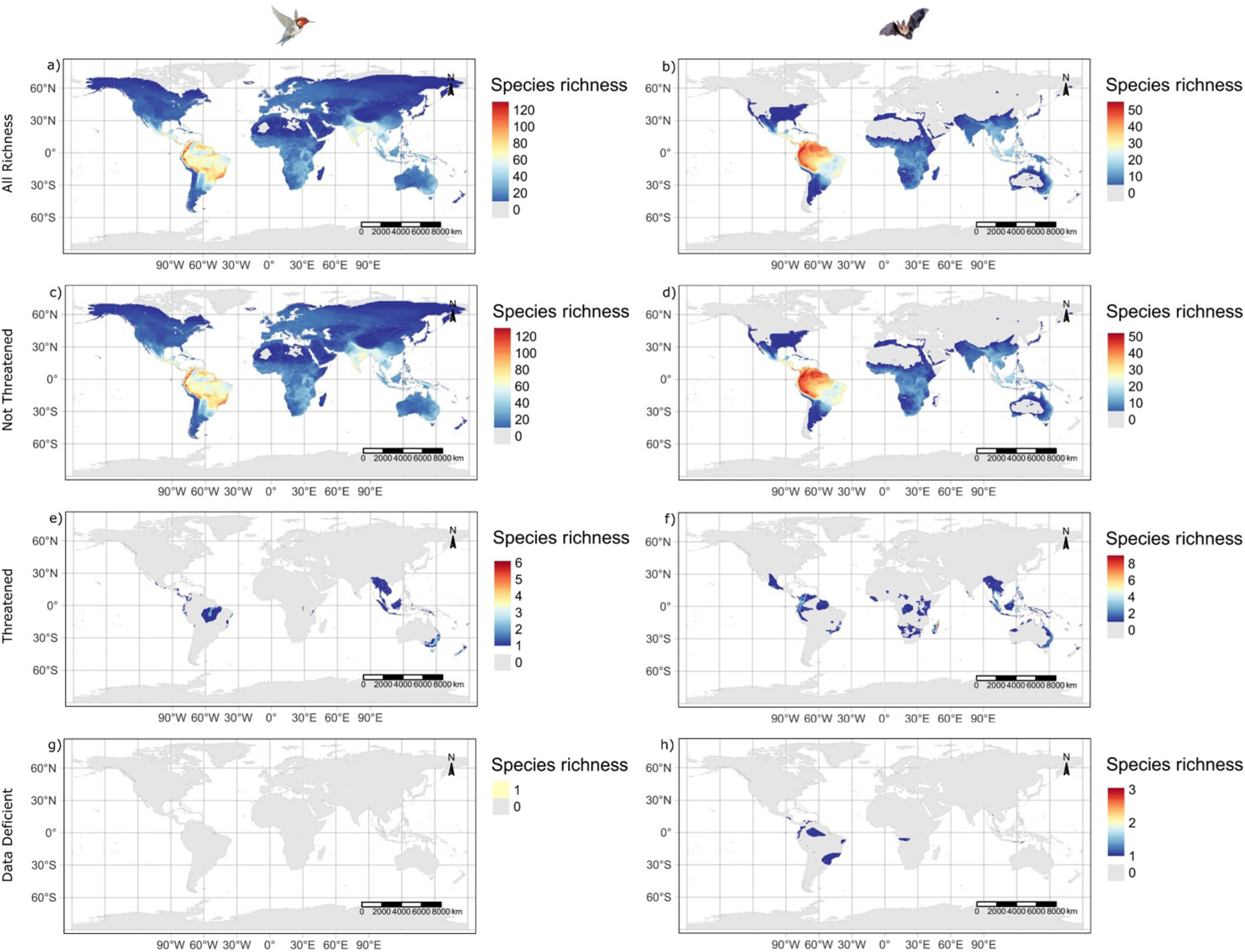
Spatial richness patterns of (a, c, e, g) bird and (b, d, f, h) mammal pollinators of the world. (a, b) All bird and mammal pollinators, (c, d) species not-threatened, (e, f) species threatened and (g, h) data deficient species. A spatial resolution of 0.5-degree latitudinal and longitudinal grids (approximately 50 km) was used to create maps

Similar to the overall richness patterns, our findings indicated that hotspots for all not-threatened bird pollinators were primarily concentrated in the tropical Andes and the Atlantic Forest regions (see Figure 1 c). Conversely, hotspots for all not-threatened mammal pollinators were observed in the north of South America (see Figure 1d). We identified Colombia and the Hawaiian Islands as hotspots for threatened bird pollinators (see Figure 1 e), while Madagascar emerged as the primary hotspot for threatened mammal pollinators (see Figure 1f). While accounting the proportion of threatened pollinators, we found Mauritius as the hotspot for bird pollinators, and Madagascar, Central Africa, Australia and New Zealand as the hotspots for mammal pollinators (see Figure S4). Additionally, the five threatened reptile pollinators were predominantly found in various islands, including Mallorca in Spain (*Podarcis lilfordi*), New Zealand (*Toropuku stephensi*), Reunion Island in Mauritius (*Phelsuma borbonica*), Socotra in Yemen (*Hemidactylus dracaenacolus*), and the Turks and Caicos Islands in Central America (*Cyclura carinata*) (see Figure 2c).

**Figure 2.**
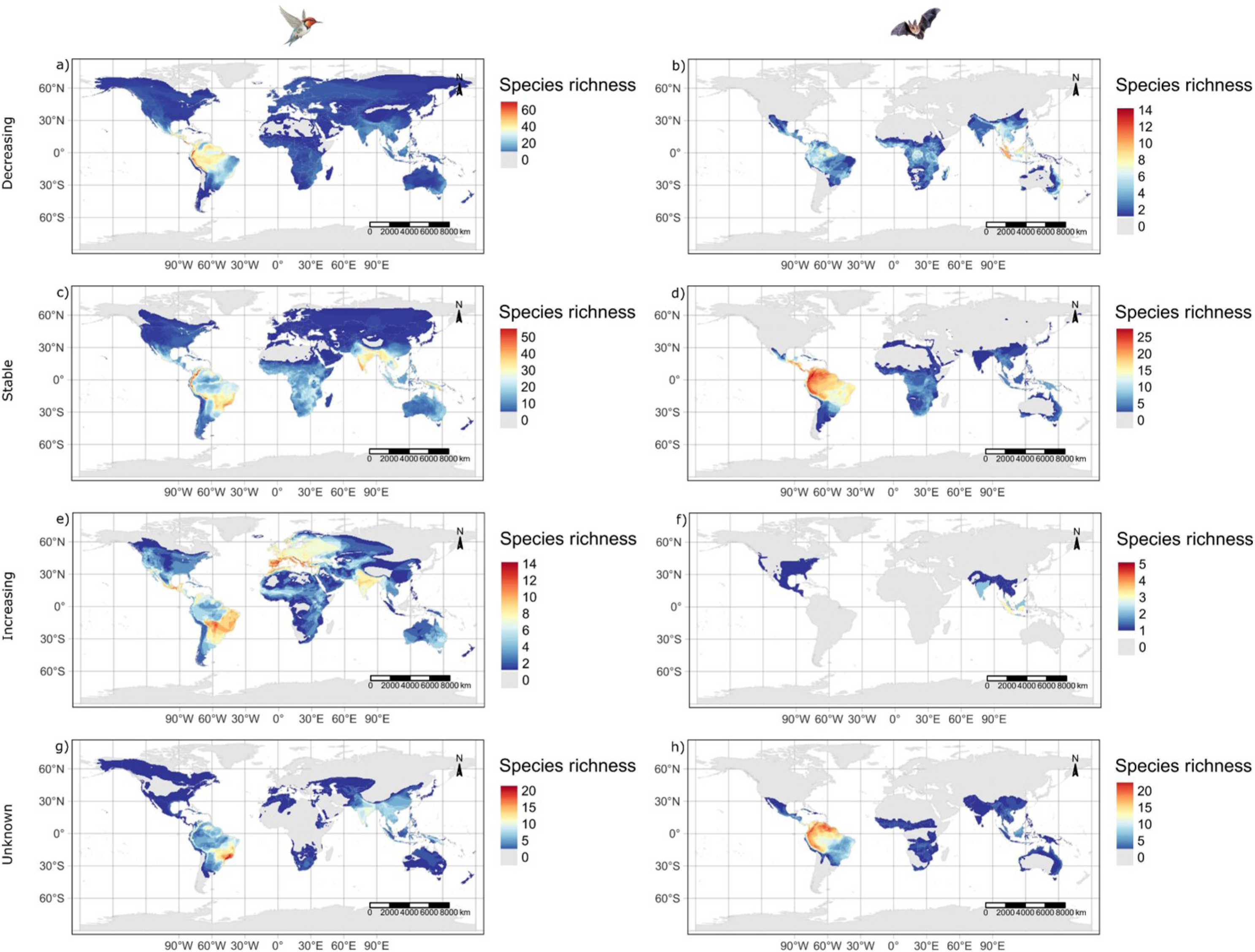
Spatial richness patterns of (a, c, e, g) bird and (b, d, f, h) mammal pollinators’ population trends. Species are organized as per the IUCN criteria. (a, b) Species with decreasing populations, (c, d) stable, (e, f) increasing and (g, h) unknown-populations. A spatial resolution of 0.5-degree latitudinal and longitudinal grids (approximately 50 km) was used to create maps

Furthermore, our results revealed two data deficient bird pollinators, notably found in Bolivia (*Discosura letitiae*) and the islands of Papua New Guinea (*Myzomela albigula*; see Figure 1g). Conversely, the hotspot for data deficient mammals was observed in the Andes region of Ecuador (see Figure 1h). But we have not mapped the data deficiency for reptile pollinators (*Pseudocordylus subviridis*).

### Spatial patterns of population trend

Our analyses investigating population trends revealed distinct patterns for bird and mammal pollinators. Hotspots of population decrease for bird pollinators were observed in the tropical Andes, Costa Rica, and Panama regions (see Figure 2a). Conversely, for mammal pollinators, this was predominantly located in Indonesian islands and eastern regions of Australia (see Figure 2b). But reptile pollinators are decreasing in western region of Mexico (see Figure S2b). Similarly, hotspots of stable populations for bird pollinators were recorded in the Andes and Atlantic Forest regions of South America and across the Indian subcontinent (see Figure 2c). Interestingly, South America also formed the hottest hotspot for mammal pollinators who have stable population trends (see Figure 2d). However, the central South America found to be the hotspots of bird pollinators whose population is increasing (see Figure 2e), but for mammal pollinators it was primarily located in the islands of Indonesia (see Figure 2f). Regarding the species with unknown population trend, hotspots for bird pollinators peaked in the Atlantic Forest regions of Brazil (see Figure 2g), whereas for mammal pollinators, it predominantly occurred in the tropical regions of South America (see Figure 2h).

### Patterns of spatial correlation of data deficient and threatened pollinators

Our analysis aimed to explore spatial correlation between data deficient and threatened bird pollinators revealed a weak correlation between them (Tjøstheim’s coefficient = 0.355 ± 0.0003). However, no correlation was observed between data deficient and threatened mammal pollinators (Tjøstheim’s coefficient = 0.056 ± 0.0001), nor between threatened bird and threatened mammal pollinators (Tjøstheim’s coefficient = -0.148 ± 0.0001). Similarly, our results indicated no spatial correlation between globally threatened birds and threatened pollinator birds (Tjøstheim’s coefficient = 0.139 ± 0.0009; see Figure S5a), and globally threatened mammals and pollinator mammals (Tjøstheim’s coefficient = -0.055 ± 0.0009; see Figure S5b).

### Taxonomic patterns of Anthropogenic threat

Our result showed that both bird and mammal pollinators were predominantly threatened from agriculture (for example, agricultural expansions, residential developments, and natural system modifications) and overexploitation (for example, hunting and poaching). In contrast to this, the reptile pollinators were generally threatened from invasive species and disease rather than agriculture or overexploitation (see Table S15). Though the mammal pollinators were significantly more threatened than bird pollinators from them (see Table S15; Figure 3a). However, we did not find a significant difference in threat between bird and mammal pollinators from climate change and invasive species (see Table S15). Among the families of bird and mammal pollinators, the Trochilidae and *Pteropodidae* pollinators were largely threatened from these factors (see Table S16; Figure S6). Additionally, we found that bird, mammal, and reptile pollinators were utilised in various markets, including food and pets. A significant proportion of mammal pollinators were extensively utilised in food (35%) and medicinal (4.3%) markets compared to bird pollinators, which were significantly utilized in pet markets (58.5%; see Figure 3b & Table S17). Likewise, the reptile pollinators were mainly used in pets (28%) than the food markets (see Table S17).

**Figure 3.**
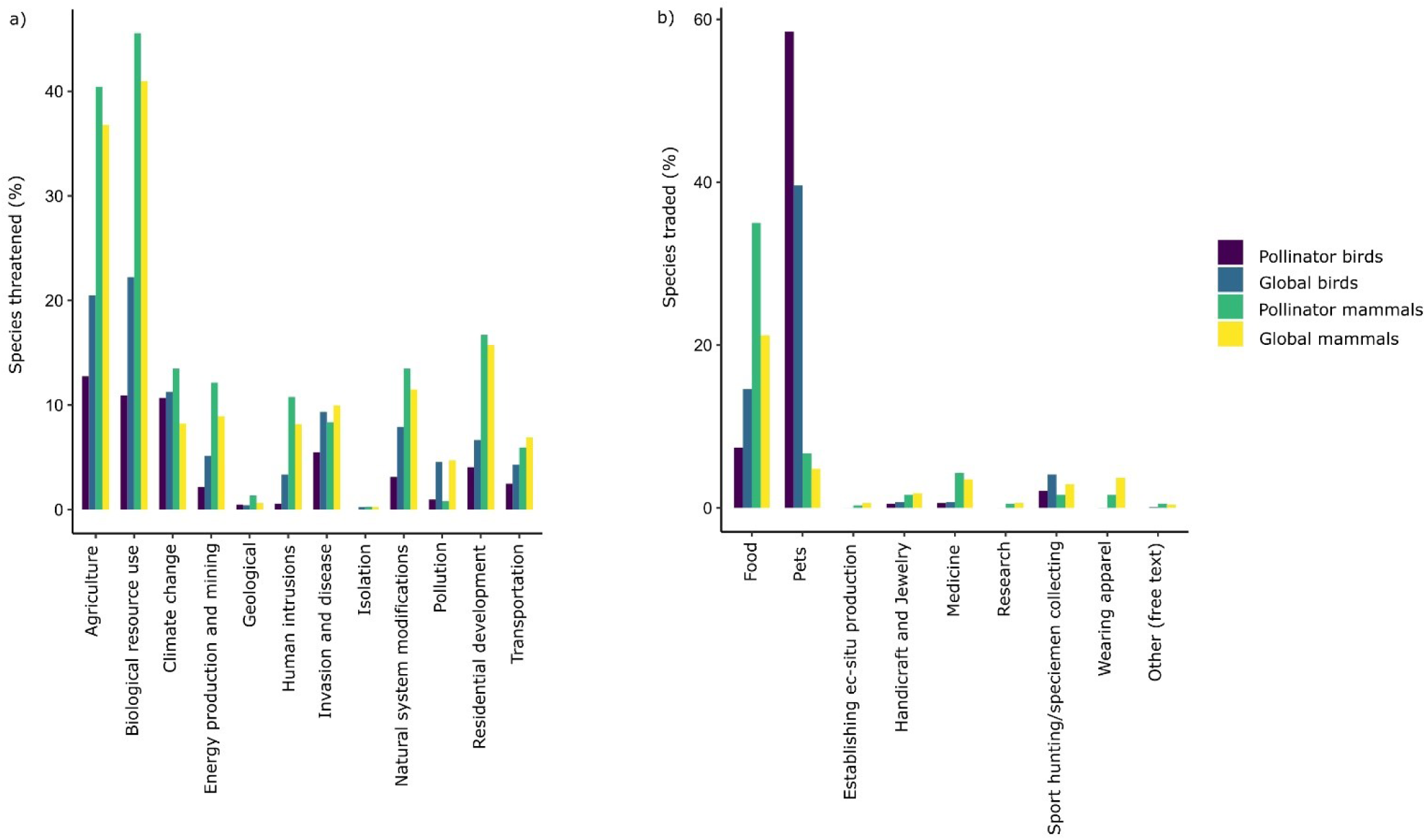
Patterns of (a) anthropogenic threat and (b) use and trade of global and pollinator bird and mammal species of the world. Some species are subject to more than one threat. The data has been extracted from the IUCN portal

### Spatial patterns of Anthropogenic threat

According to our analysis, the tropical Andes and Atlantic regions of South America were found to be hotspots where bird pollinators are threatened by habitat destruction (see Figure S7a). However, the hotspots of bird pollinators threatened from overexploitation, climate change and invasion were found to be in Australia and New Zealand (see Figure S7c, S7e & S7g). Likewise, Southeast Asia and South America have been the hotspots of habitat destruction for mammal pollinators (see Figure S7b). However, the peak hotspots of mammal pollinators threatened by overexploitation were found across Southeast Asia and parts of Africa (see Figure S7d). But the mammal pollinators threatened by climate change and invasion were found to be in eastern parts of Australia (see Figure S7f-h).

Our findings indicated that South Asia emerged as the primary hotspot for bird and mammal pollinators hunted for food markets (see Figure S8a & b). Conversely, South America and Madagascar were identified as the predominant hotspots for species utilized within pet markets (see Figure S8c & d).

### Phylogenetic pattern of extinction risk and population decline

We detected a phylogenetic signal in threat and population decline of bird and mammal pollinators without data deficient and unknown population trend species and by considering them as data-sufficient species, i.e., threatened or not-threatened (see Table S18).

## Discussion

Our estimation using the IUCN data showed an asymmetrical pattern on the extinction risk of world’s vertebrate pollinators where mammal pollinators are currently facing the greatest threat (29%; range 27.5-32%) compared to bird (8.3%; range 8.3-8.5%) and reptile (13%; range 12.5-15%) pollinators. This is consistent with the pattern of threat reported for global vertebrate species where mammals (26%) are more threatened than bird (12%) and reptile (21%) species (IUCN, 2024). Similarly, a recent study in India has identified a similar trend of highest threat where they report 28% of mammal pollinators are currently threatened with extinction risk (Kallivalappil et al. 2024). Our findings specifically highlight the need of investing more conservation effort to Critically Endangered vertebrate pollinators because no Critically Endangered bird pollinators being qualified to move to a lower category in the last 30 years (see Table S19). This is true with the mammal and reptile pollinators’ status except for the mammalian pollinator *Phyllonycteris aphylla* (see Tables S20 & S21). This aligns with the previous findings which documented the uplist of 18 bird and 13 mammalian pollinator species to a higher category of Red List (for example, Endangered to Critically Endangered) due to deterioration in their status during a period of 1988-2012 and 1996-2008 (Regan et al. 2015). Importantly, we identify insufficiently updated Red List assessment for some of our threatened pollinators (for example, *Burramys parvus* last updated in 2008; see Tables S19, S20 & S21). This insufficient monitoring can hamper their proper conservation plannings as regular monitoring is important for decision making, informing policy and to implement effective conservation programmes (Regan et al. 2015).

Our understanding of the status of current threat in vertebrate pollinators can be affected by the lack of proper Red List status for some pollinators. Therefore, our assessment will be slightly altered if the conservation of those species is impacted. For example, we found 2 birds, 16 mammals and 1 reptile pollinator in our dataset as data deficient and some of them belong to data deficient category since the past one to three decades (see Tables S19, S20 & S21). Interestingly, most of these pollinator’s population trend is currently unknown (birds = 2, mammals = 15, reptiles = 1), therefore, field studies are urgently needed for these species to identify their demographic trends because the species demographic data is an important tool for designing species conservation policies and, thus, to develop extinction risk assessments (Conde et al. 2019).

In terms of the population trends, a significant proportion of populations of bird (46%; range 40-52%), mammal (55%; range 41-67%) and reptile (14%; range 10-38%) pollinators are decreasing. Additionally, employing the IUCN data, our results suggested that 84% of the threatened bird (VU, EN, CR), 90% of the threatened mammal, and 75% of the threatened reptile pollinators’ population is currently declining (see Table S22). Such a high decline in the abundance of threatened pollinators may have implications prior to their local or global extinction (Collen et al. 2011, Ceballos et al. 2017, Isbell et al. 2023). The loss of population is a conservation concern for the threatened pollinators especially the pollinators like bats are principally susceptible to population decline because of their low reproductive rate and high age of maturity, lowering their chances of immediate population recovery (Jones et al. 2003). Therefore, advancing the understanding of the population trends of pollinators benefits proactive conservation actions and allows for targeting of both populations and species that are at immediate risk (Collen et al. 2011). Similarly, the decrease in population size of common and widespread bird (42%) and mammal (37%; see Table S22) pollinators indicate implications for ecosystem integrity and need for incorporating measures to slow down population declines of common pollinator species (Bowler et al. 2019, Rosenberg et al. 2019, Finn et al. 2023, Rigal et al. 2023).

### Extinction risk and population decrease in endemic pollinators

Further, the worrisome status of threat of endemic bird (22.5%; range 22.4-22.8%), mammal (58%; range 55.5-60%) and reptile (13; range 13-13%) pollinators suggesting that their biodiversity is on the verge of extinction. The disproportionate threat of pollinators, especially mammalian pollinators, is probably due to narrow-range size and the exposure to human threats (Craig et al. 1994, Kitahara and Sei 2001, Ceballos and Ehrlich 2002, Horner-Devine et al. 2003, Epstein et al. 2009, Mickleburgh et al. 2009, Sheherazade and Tsang 2015). Many mammal pollinators, especially bats, are mostly endemic to the Pacific and Indian Ocean islands where they are more vulnerable to exploitation pressures, invasion, demographic collapse, and environmental stochastic events (Craig et al. 1994, Pierson et al. 1996, Mickleburgh et al. 2009, Sheherazade and Tsang 2015, Mildenstein et al. 2016, Ripple et al. 2016, Vincenot et al. 2017, Newbold et al. 2018, Low et al. 2021, Kingston et al. 2023). Worryingly, more than three-fourth of the threatened endemic vertebrate pollinators’ population is currently decreasing (see Table S23). This is currently threatening the mutualistic interactions and plant reproductions in several islands of Asia-Pacific, Australia and New Zealand (Paton 1993, Vaughton 1996, Cox and Elmqvist 2000, Benning et al. 2002, Celebrezze and Paton 2004, Kelly et al. 2006, Mortensen et al. 2008, Kaiser-Bunbury et al. 2010, Anderson et al. 2011, Iles and Kelly 2014, Bissessur et al. 2019). The absence of conservation efforts to mitigate further population decline in these pollinators can impact ecosystem functioning and homogenization of the communities because they have unique combinations of functional traits that contribute to the community uniqueness (Ceballos and Ehrlich 2002, Gaston and Fuller 2008, Burlakova et al. 2011, Mouillot et al. 2013, Dirzo et al. 2014, Young et al. 2016, Newbold et al. 2018, Kidane et al. 2019, Crabot et al. 2021).

### Taxonomic variations in threat and population decrease

The taxonomic difference in threat and population decrease of bird and mammal pollinator groups and between their families varied remarkably. For example, the families such as Fringillidae, Psittacidae and Zosteropidae of birds and Pteropodidae of mammal pollinators are more threatened than their counterparts (see Table 1). This suggests that pollinators of some families are more sensitive to different threats than others, a similar pattern noticed with other vertebrate species (Bennett and Owens 1997, Ducatez and Shine 2017). But this has not been tested, therefore further research to investigate the variations in threat between taxonomic families is an urgent work for establishing global conservation priorities, identifying regions where species are particularly vulnerable to threats from humans and forecasting how threatened species maybe distributed in a changing environment (Howard et al. 2020).

### Spatial patterns of pollinators richness

Using the IUCN data, our result is able to identify knowledge gaps about various spatial patterns of vertebrate pollinators biodiversity. The higher species biodiversity of bird and mammal pollinators across the tropical parts of the world compared to other parts suggests the importance of these species to maintain the stability of tropical biodiversity (Heithaus et al. 1975, Stiles 1977, Gould 1978, Janson et al. 1981, Yumoto et al. 1997, Sazima et al. 1999, Muchhala and Jarrin-V 2002, Corlett 2004, Machado and Lopes 2004, Fleming 2005, Fleming and Muchhala 2008, Ratto et al. 2018, Vizentin-Bugoni et al. 2018). Particularly, many vertebrate pollinators across these regions evolved to establish a specialized mutual relationship with their plants because of greater phenotypic variations (i.e., morphological matching between bill of pollinators and flowers) and richness of vertebrate pollinated plant species along with the climatic stability within and between seasons which determines consistent year-round food availability to survive (Fleming 2005, Abrahamczyk and Kessler 2010, 2015, Higgins et al. 2016, Zanata et al. 2017). Though we notice the diversity of vertebrate pollinators in temperate regions of, for example, eastern parts of Australia. This is possibly due to their broader dietary preferences (such as nectar, fruits, insects, honeydew) allowing them to occupy such regions (Pyke 1980, Higgins et al. 2016).

Additionally, the pattern of higher richness of both bird and mammal pollinators in South America relative to other regions possibly driven by the predictability of resource in space and time (Waser et al. 1996, Ollerton et al. 2006, Fleming and Muchhala 2008, Zanata et al. 2017, Rodriguez-Flores et al. 2019), but we have not tested this. Studies proposed that the phylogenetic history and spatio-temporal predictability of flower resources (higher diversity of potentially vertebrate-pollinated plants) are largely responsible for hemispheric and regional differences in the structure of vertebrate pollinator assemblages (Fleming 2005, Fleming and Muchhala 2008, Zanata et al. 2017, Dalsgaard et al. 2021). Interestingly, the higher resource diversity and spatio-temporal predictability in South America facilitated the spread of many nectar-feeding birds and bats into a variety of relatively specialized niches compared to reduced resource diversity and spatio-temporal predictability regions like Asia and Australia (Sakai 2001, Fleming 2005, Fleming and Muchhala 2008). In contrast, we notice most reptile pollinators are important on various island ecosystems. This is due to the high densities and lower predation risk on islands that often facilitate them to expand the dietary niche to nectar and pollen thus act as pollinators (Olesen and Valido 2003).

### Spatial patterns of threat and population decrease

Our investigation to explore the spatial congregation of threatened pollinators biodiversity showed varying patterns. The spatial difference in threat between the global bird (Himalaya and southeast Asia) and mammal (Southeast Asia) species and pollinator bird (Colombia, Hawaiian Islands, Australia and Southeast Asia) and mammal (Madagascar) species emphasizes the need for more species-specific conservation actions to prevent extinction of the threatened vertebrate pollinators (Roll et al. 2017, Cox et al. 2022). Additionally, the decline of population of pollinators suggests a more comprehensive population monitoring programme is crucial for informing conservation actions and reassessing the status of species (Stephenson et al. 2022). A change in pollinators population critically affect the genetic and functional diversity of both pollinators and plants which consequently result in lower the quality of pollination services, gene flow and recruitment of new plants (Mortensen et al. 2008, Anderson et al. 2011, Cianciaruso et al. 2013, Bustamante et al. 2016). Hence, population monitoring is an urgent task because many vertebrate pollinators’ distribution spans in poor, high-biodiversity countries across the tropics, therefore the extinction or loss of pollinators will largely collapse the biodiversity, ecosystem services and the economy of people in this region (Bumrungsri et al. 2008, Bumrungsri et al. 2009, Kunz et al. 2011, Bumrungsri et al. 2013, Whelan et al. 2015, Trejo-Salazar et al. 2016, Aziz et al. 2017, Stephenson et al. 2022). The decline of reptile pollinators principally impacts several endemic plants on islands (Olesen and Valido 2003, García and Vasconcelos 2017, Romero-Egea et al. 2023).

The dominant threat to vertebrate pollinators across these regions is mainly driven by habitat destruction for agriculture and developments than climate change and overexploitation (see Table S15). This is consistent with the previous findings (Regan et al. 2015, Kallivalappil et al. 2024), and more it reflects the general patterns observed for the global bird and mammal species (Hoffmann et al. 2010, Hogue and Breon 2022). Existing evidence shows that the response of bird pollinators varies to habitat destruction. For example, the urbanization and fragmentation of habitat reduced the functional diversity and population growth rate of sunbird pollinators in South Africa and Tanzania (Stouffer and Bierregaard Jr 1995, Korfanta et al. 2012, Pauw and Louw 2012, Kormann et al. 2016). It is believed that hummingbirds are resilient to habitat disturbance (Borgella Jr et al. 2001, Morrison and Mendenhall 2020). But the accumulating evidence suggests that habitat destruction influences the functional composition of hummingbird pollinators’ communities and thus affects plant mating quality and fitness (Borgella Jr et al. 2001, Hadley et al. 2014, Hadley et al. 2018, Torres-Vanegas et al. 2021). Worryingly, this is detrimental to the specialized hummingbirds by restricting their mobility between forest-patches (Kormann et al. 2016, Morrison and Mendenhall 2020, Torres-Vanegas et al. 2021). As the relationship between these specialist hummingbird pollinators tends to be more closely interconnected with their native plant species (Leimberger et al. 2022, Barreto et al. 2024). There is evidence that the decline in reproduction of *Heliconia* is likely driven by the restricted movement of specialist hummingbirds between forest fragments (Hadley et al. 2014).

The impact of habitat destruction is obviously not unique to bird pollinators, but growing evidence shows that it is increasingly detrimental to mammal pollinators (Mickleburgh et al. 2002, Tarnaud and Simmen 2002, Kingston 2010, Volampeno et al. 2013, Aziz et al. 2021). Despite some degree of tolerance to habitat modifications, the cyclone and agriculture related activities negatively affect the abundance and richness of bat pollinators across the world (Craig et al. 1994, Pierson et al. 1996, McConkey et al. 2004, Meyer et al. 2008, Meyer and Kalko 2008, Voigt and Kingston 2016, Rocha et al. 2017, Webala et al. 2019, Lavery et al. 2020). Such uncontrolled land clearings and modifications further reduce their population by limiting the opportunity for foraging and roosting. Therefore, it emphasizes that the conservation of vertebrate pollinators can only be achieved through the protection of large-scale habitats. Failure to protect habitat may be the reason why we have failed repeatedly to achieve biodiversity targets set under the Convention on Biological Diversity; therefore, habitat destruction regulation must be at the forefront of the global environmental priority programmes, particularly conservation hotspots must be prioritized (Tittensor et al. 2014, Hogue and Breon 2022).

Climate change is the second most prevalent threat we observe for bird pollinators followed by invasive species impact. It disproportionately changes the structure and function of hummingbird communities and populations by shifting ranges altitudinally upwards, latitudinally northward and limiting the opportunity to expand further (Waser 1976, Lara et al. 2012, McKinney et al. 2012, Şekercioğlu et al. 2012, Courter 2017). The severity of climate change will be more on montane endemic bird pollinators where they have limited capacity to shift ranges (Şekercioğlu et al. 2012). This ultimately threatens the plant reproductions and mutualistic interactions by reducing the range overlap between plants and their pollinators (Adedoja et al. 2024). For example, the phenological mismatch between the flowering plants and arrival of migrant hummingbirds (*Selasphorus platycercus*) can potentially affect the reproduction of hummingbirds and plants, leading to population decline of both groups (Waser 1976, McKinney et al. 2012). In the Hawaiian Islands, climate change along with the invasive disease (avian pox and malaria) causing mosquitoes makes the survival of honeycreeper (Drepanidinae) pollinators difficult. In many islands they have already shifted to the higher elevations where they are considered the last refugia for these pollinators (Warner 1968, Banko et al. 2002, Benning et al. 2002).

The higher threat of mammal pollinators from overexploitation is majorly attributed to the widespread hunting of fruit-bats (*Pteropus* spp.) for consumption, medicinal properties, and sport. Bushmeat hunting is increasingly recognized as a major conservation threat to fruit-bat pollinators across Southeast Asia and West and Central Africa (Epstein et al. 2009, Mickleburgh et al. 2009, Mildenstein et al. 2016, Ripple et al. 2016), and to the primate pollinators of South America (Bogoni et al. 2020). Although all *Pteropus* species have been listed in appendix I or II of CITES (Convention on International Trade in Endangered Species) since 1989, thus prohibiting the trade of legally hunted bats in international markets (Vincenot et al. 2017). But local hunting, legal or illegal mass culls, legal persecution and relaxed law enforcement make the biggest challenge for bat pollinators conservation (Wiles 1987, Jones et al. 2009, Florens et al. 2017, Vincenot et al. 2017, Sheherazade and Tsang 2018, Florens and Baider 2019, Frick et al. 2020, Tackett et al. 2022, Kingston et al. 2023). As such increasing pressures on bats fundamentally drive population declines and reduce their role as keystone species which often disrupt the ecological interactions that constitute this keystone role (Florens et al. 2017, Ratto et al. 2018). We found invasive species to be the greatest threat to reptile pollinators. This is due to the introduction of invasive species on islands (Cooper Jr and Pérez-Mellado 2012, Buckland et al. 2014).

Using the IUCN data, we found commercial trade of various vertebrate pollinators including hummingbird species’ illegal trade in the markets of Mexico and USA (Gómez Álvarez and Reyes Gómez 2010, Ebersole 2018). While unknown the magnitude of harvest, it is documented that the populations of some hummingbird species are increasingly declining (Sauer et al. 1966, Sauer et al. 2013, English et al. 2021). The increased demand for trades and unsustainable illegal harvesting in Southeast Asia, particularly in Indonesia, are identified as the greatest threat to many sunbirds and white-eyes pollinators in this region (Su et al. 2014, Eaton et al. 2015, Su et al. 2015, Chng et al. 2018a, Chng et al. 2018b, Leupen et al. 2022a, Leupen et al. 2022b). Notably, the heavy trade of endemic bird pollinators impacted the steep decline of local populations. For example, a recent study shows that 80% of population decline in a threatened white-eye species (*Zosterops flavus*) in the last 10 years because of heavy trade in Java, Indonesia, (van Balen et al. 2023). Therefore, the use of novel methods like application of bioacoustics and deep learning (for identifying cryptic species) will be critical for monitoring and tackling the bird trade markets, protecting threatened species, and enforcing wildlife laws (Su et al. 2024). Despite the diversity bats rarely trade/consume in tropical South America, but the uncontrolled internal commercial trade is a pressing conservation issue for fruit-bat pollinators in several low-income countries of Africa, Asia and many Indian and Pacific Ocean islands where they often trade as bat bushmeat from 1 to 500 metric tons annually (Mickleburgh et al. 2009, Sheherazade and Tsang 2015, Tanalgo et al. 2023). Therefore, incorporating options such as enhancing domestic meat, generating opportunities for additional source of income, engage stakeholders in pollinators conservation, increasing investments in the rural economy, and improve laws and strengthen enforcement of existing protective legislation are essential for effective conservation of vertebrate pollinators world-wide (Sheherazade and Tsang 2015, Chapron et al. 2017, Vincenot et al. 2017, Kingston et al. 2023, Tanalgo et al. 2023).

### Spatial correlation of data deficient and threatened pollinators biodiversity

We are able to highlight a weak spatial correlation between data deficient and threatened bird pollinators. A recent study has found that many data deficient species are in fact more threatened than the data-sufficient species (Borgelt et al. 2022). It is therefore reasonable to believe that the data deficient bird pollinators are experiencing threats in these regions. However, we could not find such a relationship with mammal pollinators, suggesting less likelihood of experiencing threat in data deficient mammal pollinators. However, this relationship is scale dependent (McPherson et al. 2006, Keil et al. 2011).

### Phylogenetic signal in threat and population decline

Besides anthropogenic threats, a striking finding in our study was the presence of phylogenetic signal in threat and population decline of both bird and mammal pollinators, which corroborates with the previous findings (Bennett and Owens 1997, Purvis et al. 2000b, Wang et al. 2018). This indicates that both threatened bird and mammal pollinators are evolutionarily predisposed to extinction risk and population decline, hence, such non-random nature of extinction risk can reduce vertebrate pollinators’ biodiversity much more than the random nature of extinction (Bennett and Owens 1997, Purvis et al. 2000a, Jones et al. 2003). Therefore, these threatened clades need immediate conservation attention otherwise it causes disproportionate loss of vertebrate pollinators biodiversity (Jetz et al. 2012, Nunes et al. 2015, Wang et al. 2018).

## Conclusion

Using a newly built dataset and the IUCN information, we show the pattern of extinction risk and population decline of vertebrate pollinators world-wide. We show elevated extinction risk for mammal pollinators relative to birds and reptiles. The threat to birds and mammals is mainly driven by both external and internal factors. The external drivers of extinction are mostly related to extrinsic anthropogenic factors such as agricultural practice and overexploitation, particularly the type of human uses with birds being subdued to the pet market and mammals being hunted for bushmeat trade. The internal threat shows the pattern of extinction risk in related species. It is interesting to explore further research on ecological and evolutionary instinct of pollinators to threat, and how trade related hunting impacts phylogenetically related species and their response to changing climate. The hotspots of threatened vertebrate pollinators in low-income and densely populated countries of Africa and Asia are a pressing challenge for vertebrate pollinators conservation due to several socioeconomic factors and the increasing magnitude of different threats. Therefore, formulating the novel policies, land-use regulation and legislation on hunting must be implemented along with promoting alternative opportunities such as wildlife watching and ecotourism in these countries. Our result further showed that many common and wide-spread pollinators including many island endemics are also declining. This highlights that special attention must be given to the endemic and island vertebrate pollinators accounting their vulnerability to anthropogenic and climate pressure. The insufficient Red List status and population trend data for some species may underestimate the actual breadth of pollinators’ loss; therefore, field work and periodic monitoring should be encouraged for innovative conservation plans and to mitigate extinction of vertebrate pollinators across the planet.

## Supporting information

https://zenodo.org/uploads/14515205

## Notes

https://zenodo.org/uploads/14515205

### Competing Interest Statement

The authors have declared no competing interest.

