## Supplementary material for "The decline of global pollinator biodiversity in the Anthropocene": https://zenodo.org/uploads/14515205

### *Tables*

**Table: S1 Vertebrate pollinator selection criteria employed by Regan et al. (2015) for bird and mammal pollinators, and the criteria developed for reptile species to be considered as pollinators for this study.**

| | Group | Criteria |
|---|---|---|
| Regan et al. (2015) | Bird | Included all species in the families Coerebidae, Meliphagidae, Mohoidae, Nectariniidae, Promeropidae, Trochilidae, and Zosteropidae, plus selected species of other families including Fringillidae, Icteridae, Psittacidae, and Thraupidae, drawing on descriptions of foraging behaviour and diet. |
| | Mammal | Included species that have been regularly observed sucking or licking flowers' nectar, or carrying pollen load on fur, or in the case of bats, those that are predicted to be pollinators based on their tongue morphology. |
| Kallivalappil et al | Reptile | We defined a pollinator species based on descriptions of foraging behaviour such as contact with reproductive structures of flowers with their throat or snout, and diet or reported carry pollens as per the data suggestion or experiments show fruit/seed set formation followed by animals visit. |

**Table S2 Keywords used to retrieve studies from the Web of Science and Google Scholar databases. Each key word and its corresponding retrieved and selected study (numbers) is given with them**

| **Web of Science (search: 1985 – 2022)** | | | | | |
|---|---|---|---|---|---|
| **Keywords** | **Retrieved** | **Selected** | **Keywords** | **Retrieved** | **Selected** |
| Vertebrate polli* | 525 | 88 | lizard flowe* | 132 | 29 |
| Bird polli* | 2,019 | 426 | bird plant reproduct* | 1,843 | 283 |
| Avian polli* | 246 | 35 | bat plant reproduct* | 376 | 131 |
| Passerine bird polli* | 106 | 50 | vertebrate plant interact* | 1,677 | 129 |
| Mammal polli* | 532 | 92 | bird plant interact* | 2,738 | 183 |
| Bat polli* | 1,005 | 183 | mammal plant interact* | 2,363 | 44 |
| Reptile polli* | 46 | 2 | bat plant interact* | 670 | 79 |
| Lizard polli* | 108 | 30 | reptile plant interact* | 135 | 3 |
| ornithoph* | 491 | 76 | lizard plant interact* | 203 | 19 |
| vertebrate flowe* | 672 | 73 | vertebrate flowe* reprod* | 140 | 19 |
| bird flowe* | 2,463 | 274 | bird flowe* reprod* | 559 | 227 |
| mammal flowe* | 868 | 95 | mammal flowe* reprod* | 192 | 20 |
| bat flowe* | 1235 | 165 | bat flowe* reprod* | 246 | 68 |
| reptile flowe* | 87 | 3 | | | |
| **Google Scholar (search: until 31 December 2022)** | | | | | |
| **Keywords** | **Retrieved** | **Selected** | **Keywords** | **Retrieved** | **Selected** |
| Vertebrate+pollination | 300 | 182 | Mammal+pollination | 300 | 198 |
| Bird+pollination | 300 | 273 | Bat+pollination | 300 | 277 |
| Avian+pollination | 300 | 128 | Reptile+pollination | 300 | 22 |
| Passerine+bird+pollination | 300 | 177 | Lizard+pollination | 300 | 54 |
| **Other Sources** | | **Selected** | | | |
| Singh, 1929; Ali, 1931; McCann, 1931, 1933; Faegri and Pijl, 1961; Davidar, 1985; Raju 1990, 2008; Proctor et al. 1996; Nicodemus et al. 2006 | | 10 | | | |

**Table S3 Bird and mammal pollinator species recorded as extinct in Regan et al. (2015) dataset according to the IUCN Red List status 2012 (birds) and 2008 (mammals).**

| Birds | Red List 2012 | Red List 2022 | Mammals | Red List 2008 | Red List 2022 |
|---|---|---|---|---|---|
| *Anthornis melanocephala* | Extinct | Extinct | *Pteropus tokudae* | Extinct | Extinct |
| *Chaetoptila angustipluma* | Extinct | Extinct | | | |
| *Chlorostilbon bracei* | Extinct | Extinct | | | |
| *Chlorostilbon elegans* | Extinct | Extinct | | | |
| *Ciridops anna* | Extinct | Extinct | | | |
| *Drepanis funerea* | Extinct | Extinct | | | |
| *Drepanis pacifica* | Extinct | Extinct | | | |
| *Hemignathus ellisianus* | Extinct | Extinct | | | |
| *Hemignathus lucidus* | Extinct | Extinct | | | |
| *Hemignathus obscurus* | Extinct | Extinct | | | |
| *Hemignathus sagittirostris* | Extinct | Extinct | | | |
| *Moho apicalis* | Extinct | Extinct | | | |
| *Moho bishopi* | Extinct | Extinct | | | |
| *Moho braccatus* | Extinct | Extinct | | | |
| *Moho nobilis* | Extinct | Extinct | | | |
| *Zosterops conspicillatus* | Endangered | Extinct | | | |
| *Zosterops strenuous* | Extinct | Extinct | | | |

**Table S4 The IUCN shapefiles and corresponding codes of each pollinator's shapefile. For the code Presence: 1 = extant, 2 = probably extant, 3 = possibly extant, 4 = possibly extinct, 5 = extinct (post 1,500), 6 = presence uncertain. For the code Origin: 1 = native, 2 = reintroduced, 3 = introduced, 4 = vagrant, 5 = origin uncertain, 6 = assisted colonisation. For the code Seasonality: 1 = resident, 2 = breeding season, 3 = non-breeding season, 4 = passage, 5 = seasonal occurrence uncertain. The data has been collected from the IUCN portal.**

| Groups | Species | Presence | Origin | Seasonality |
|---|---|---|---|---|
| **Bird** | *Discosura letitiae* | 3 | 1 | 1 |
| | *Charmosyna diadema* | 4 | 1 | 1 |
| | *Eriocnemis godini* | 4 | 1 | 1 |
| | *Psittirostra psittacea* | 4 | 1 | 1 |
| **Mammal** | *Mystacina robusta* | 4 | 1 | 1 |
| | *Pteropus melanotus* | 6 | 1 | 1 |
| **Reptile** | *Cnemidophorus murinus* | 1 | 5 | 1 |

**Table S5 The summary statistics and parameters of proportion and Wilcoxon Man-Whiteny tests which used to test the difference in extinction risk between bird and mammal pollinators. The numbers correspond to the Threatened (VU+EN+CR) and Not Threatened (LC+NT) indicate the number of species belong to that category. The species with DD status was omitted from the analysis**

| | | | | Bird | | Mammal | |
|---|---|---|---|---|---|---|---|
| | $X^2$ | df | p-value | Threatened | Not threatened | Threatened | Not threatened |
| **Proportion Test** | 101.57 | 1 | < 2.2e-16 | 104 | 1,253 | 102 | 355 |
| **Man-Whiteny Test** | W | | p-value | | | | |
| | 267851 | | < 2.2e-16 | | | | |

**Table S6 The summary statistics and parameters of proportion test which used to test the difference in extinction risk between endemic and widespread bird pollinators. The numbers correspond to the Threatened (VU+EN+CR) and Not Threatened (LC+NT) indicate the number of species belong to that category. The species with DD status was omitted from the analysis**

| | | | | Endemic | | Widespread | |
|---|---|---|---|---|---|---|---|
| | $X^2$ | df | p-value | Threatened | Not Threatened | Threatened | Not Threatened |
| **Proportion Test** | 195.14 | 1 | <2.2e-16 | 93 | 338 | 11 | 811 |

**Table S7 The summary statistics and parameters of Fisher's Exact test which used to test the difference in extinction risk between endemic and widespread mammal pollinators. The numbers correspond to the Threatened (VU+EN+CR) and Not Threatened (LC+NT) indicate the number of species belong to that category. The species with DD status was omitted from the analysis**

| | | | Endemic | | Widespread | |
|---|---|---|---|---|---|---|
| | odds ratio | p-value | Threatened | Not Threatened | Threatened | Not Threatened |
| Fisher's Exact Test | 6.440 | 4.102e-13 | 56 | 40 | 46 | 213 |

**Table S8 Summary statistics of Spearman Correlation test that used to test the relationship between the family size (number of species per family) and the number of species threatened/decreasing for the families of bird pollinators. We used 64 bird families for the analysis. Data normality tested for all species (Total), threatened and decreasing species also reported separately**

| Normality Test | | | Spearman's Correlation | | | |
|---|---|---|---|---|---|---|
| | W | p-value | Birds | S | $R^2$ | p-value |
| Total species | 0.372 | 0.001 | | | | |
| Threatened | 0.369 | 0.001 | Threatened | 216 | 0.505 | 0.001 |
| Decreasing | 0.321 | 0.001 | Decreasing | 127 | 0.71 | 0.001 |

**Table S9 Summary statistics of Spearman Correlation test that used to test the relationship between the family size (number of species per family) and the number of species threatened/decreasing for the families of mammal pollinators. We used 37 bird families for the analysis. Data normality tested for all species (Total), threatened and decreasing species also reported separately**

| Normality Test | | | Spearman's Correlation | | | |
|---|---|---|---|---|---|---|
| | W | p-value | Mammals | S | $R^2$ | p-value |
| Total species | 0.347 | 0.001 | | | | |
| Threatened | 0.339 | 0.001 | Threatened | 350 | 0.585 | 0.001 |
| Decreasing | 0.374 | 0.001 | Decreasing | 198 | 0.765 | 0.001 |

**Table S10 The summary statistics and parameters of proportion and Wilcoxon Man-Whiteny tests which used to test the difference in population decrease between bird and mammal pollinators. The numbers correspond to the Decreasing and Not Decreasing (Stable+Increasing) indicate the number of species belong to that category. The species with Unknown-population trend was omitted from the analysis**

| | | | | Bird | | Mammal | |
|---|---|---|---|---|---|---|---|
| **Proportion Test** | $X^2$ | df | p-value | Decreasing | Not Decreasing | Decreasing | Not Decreasing |
| | 7.1802 | 1 | 0.007371 | 507 | 1,106 | 152 | 276 |
| **Man-Whiteny Test** | W | | p-value | | | | |
| | 166718 | | 0.006 | | | | |

**Table S11 Summary statistics and parameters of Proportion test which used to test the difference in population decrease between endemic and widespread bird pollinators. The numbers correspond to the Decreasing and Not Decreasing (Stable+Increasing) indicate the number of species belong to that category. The species with Unknown-population trend was omitted from the analysis**

| | | | | Endemic | | Widespread | |
|---|---|---|---|---|---|---|---|
| **Proportion Test** | $X^2$ | df | p-value | Decreasing | Not Decreasing | Decreasing | Not Decreasing |
| | 6.582 | 1 | 0.0103 | 193 | 376 | 314 | 730 |

**Table S12 Summary statistics and parameters of Proportion test which used to test the difference in population decrease between endemic and widespread mammal pollinators. The numbers correspond to the Decreasing and Not Decreasing (Stable+Increasing) indicate the number of species belong to that category. The species with Unknown-population trend was omitted from the analysis**

| | | | | Endemic | | Widespread | |
|---|---|---|---|---|---|---|---|
| **Proportion Test** | $X^2$ | df | p-value | Decreasing | Not Decreasing | Decreasing | Not Decreasing |
| | 12.123 | 1 | 0.0005 | 60 | 84 | 92 | 192 |

**Table S13 The summary statistics and parameters of proportional and Wilcoxon rank sum tests which used to test the spatial distributional difference between bird and mammal pollinators of world. The category Present in the bird and mammal groups indicates the presence of at least a single species in the grid cell. The category Absent indicates the absence of species in the cells. The category Total indicates the total number of grid cells for 0.5-degree longitude-latitude grids.**

| | **$X^2$** | **df** | **p-value** | **Grid cells** | | | | |
|---|---|---|---|---|---|---|---|---|
| | | | | **Bird** | | **Mammal** | | |
| | | | | **Present** | **Absent** | **Present** | **Absent** | **Total** |
| Proportion test | 27603 | 1 | < 2.2e-16 | 64,021 | 34,875 | 27,187 | 71,709 | 98,896 |
| | W | | p-value | | | | | |
| Wilcoxon test | 6956184845 | | < 2.2e-16 | | | | | |

**Table S14 The method of re-classification of various anthropogenic threat faced by bird and mammal pollinators. The species threatened from both hunting and logging has been classified under the category of overexploitation. The classification method has been adopted from González-Suárez et al. (2013) and Cox et al. (2022).**

| Category | IUCN main threat |
|---|---|
| Habitat destruction/alteration | 1. Residential and commercial development |
| Habitat destruction/alteration | 2. Agriculture and Aquaculture |
| Habitat destruction/alteration | 3. Energy production and mining |
| Habitat destruction/alteration | 4. Transportation and service corridors |
| Overexploitation | 5.1 Hunting and collecting terrestrial animals (or hunting/logging) |
| Habitat destruction/alteration | 5.2 Gathering terrestrial plants |
| Habitat destruction/alteration | 5.3 Logging and wood harvesting |
| Habitat destruction/alteration | 6. Human intrusion and disturbance |
| Habitat destruction/alteration | 7. Natural system modifications |
| Invasive species | 8. Invasive and other problematic species, genes and diseases |
| Habitat destruction/alteration | 9. Pollution |
| Climate change | 11. Climate change and severe weather |

NB: threat categories such as geological events and isolation were not included in our classification, instead kept them separately

González-Suárez, M., Gómez, A. & Revilla E. (2013) Which intrinsic traits predict vulnerability to extinction depends on the actual threatening processes. *Ecosphere*, **4(6)**, 76.

Cox, N., Young, B.E., Bowles, P., Fernandez, M., Marin, J., Rapacciuolo, G., Böhm, M., Brooks, T.M., Hedges, S.B., Hilton-Taylor, C. and Hoffmann, M., 2022. A global reptile assessment highlights shared conservation needs of tetrapods. Nature, 605(7909), pp.285-290.

**Table S15 Taxonomic patterns of anthropogenic threat in bird and mammal pollinators. The main classified threats (in bold) along with sub threats are given. The data has been extracted from the IUCN portal. Some species are subject to more than one threat. The threat percentage of bird, mammal and reptile pollinators derived from the total species threatened for specific variable against the sum of bird (1,255), mammal (371) and reptile (40) pollinators. The difference between bird and mammal pollinators in each threat was tested by a chi-square test using the function prop.test in R software**

| Threats | Birds | Threat % | Mammals | Threat % | Chi-Squared | Reptiles | Threat % |
|---|---|---|---|---|---|---|---|
| **Habitat destruction** | **194** | **15.46** | **182** | **49.06** | **< 0.001** | **8** | **20** |
| Agriculture | 160 | 12.75 | 150 | 40.43 | < 0.001 | 7 | 18 |
| Residential development | 51 | 4.06 | 62 | 16.71 | < 0.001 | 4 | 10 |
| Energy production and mining | 27 | 2.15 | 45 | 12.13 | < 0.001 | 0 | 0 |
| Transportation | 31 | 2.47 | 22 | 5.93 | 0.002 | 2 | 5 |
| Human intrusions | 7 | 0.56 | 40 | 10.78 | < 0.001 | 2 | 5 |
| Natural system modifications | 39 | 3.11 | 50 | 13.48 | < 0.001 | 1 | 3 |
| Pollution | 12 | 0.96 | 3 | 0.81 | 1 | 1 | 3 |
| Logging | 93 | 7.41 | 50 | 13.48 | 0.0004 | 1 | 3 |
| Gathering terrestrial plants | 2 | 0.16 | 0 | 0.00 | 1 | 1 | 3 |
| **Overexploitation** | **42** | **3.35** | **119** | **32.07** | **< 0.001** | **7** | **18** |
| Hunting | 16 | 1.7 | 37 | 9.97 | < 0.001 | 6 | 15 |
| Hunting/logging | 26 | 2.07 | 82 | 22.10 | < 0.001 | 1 | 3 |
| **Climate change** | **134** | **10.68** | **50** | **13.48** | **0.161** | **2** | **5** |
| **Invasion and disease** | **69** | **5.50** | **31** | **8.36** | **0.067** | **11** | **28** |
| Geological events | 6 | 0.48 | 5 | 1.35 | 0.151 | 0 | 0 |
| Isolation | 0 | 0.00 | 1 | 0.27 | 0.52 | 0 | 0 |

**Table S16 Patterns of anthropogenic threat in major families of bird and mammal pollinators. The families highlighted in bold represent the taxonomic class mammalians and the remaining families represent the taxonomic class birds**

| | Agriculture | Biological | Climate change | Energy production and mining | Geological events | Human intrusions | Invasion and disease | Natural system modifications | Pollution | Residential development | Transportation |
|---|---|---|---|---|---|---|---|---|---|---|---|
| Fringillidae | 6 | 5 | 7 | 0 | 1 | 1 | 8 | 1 | 0 | 2 | 0 |
| Icteridae | 5 | 2 | 2 | 0 | 1 | 0 | 5 | 2 | 1 | 2 | 2 |
| Meliphagidae | 12 | 11 | 34 | 3 | 0 | 0 | 6 | 4 | 0 | 1 | 2 |
| Nectariniidae | 14 | 15 | 9 | 1 | 0 | 1 | 1 | 4 | 0 | 1 | 1 |
| Psittacidae | 25 | 35 | 16 | 3 | 1 | 2 | 12 | 4 | 1 | 7 | 3 |
| Thraupidae | 7 | 2 | 2 | 2 | 0 | 1 | 4 | 1 | 1 | 5 | 4 |
| Trochilidae | 59 | 33 | 15 | 15 | 1 | 0 | 5 | 10 | 6 | 20 | 16 |
| Zosteropidae | 24 | 22 | 32 | 3 | 2 | 0 | 16 | 9 | 2 | 8 | 1 |
| **Phyllostomidae** | 19 | 16 | 6 | 16 | 2 | 16 | 3 | 4 | 0 | 7 | 2 |
| **Pteropodidae** | 67 | 81 | 33 | 15 | 3 | 16 | 12 | 12 | 2 | 24 | 7 |

**Table S17 Pattern of various use and trade of global and pollinator bird, mammal and reptile species. The number with each group (bird, mammal, reptile) and use and trade indicate the total number of species belong to that group and the number of species used in use and trade purposes along with percentage. Some species are subject to more than one use and trade purposes. The difference between bird and mammal pollinators in terms of use and trade was tested with a proportion test. The Chi-square value and its significance levels (*** = <0.001, ** = 0.001, NS = Not significant) were reported for each variable. The information about each species has been extracted from the IUCN portal.**

| Use and Trade | Pollinator birds (1,255) | % | Global birds (11,024) | % | Pollinator mammals (371) | % | Global mammals (5,886) | % | Chi-squared | Pollinator reptiles (40) | % |
|---|---|---|---|---|---|---|---|---|---|---|---|
| Food | 93 | 7.41 | 1610 | 14.60 | 130 | 35.04 | 1246 | 21.17 | 182.4*** | 2 | 5 |
| Pets | 734 | 58.49 | 4360 | 39.55 | 25 | 6.74 | 283 | 4.81 | 306*** | 11 | 28 |
| Medicine - human & veterinary | 7 | 0.56 | 78 | 0.71 | 16 | 4.31 | 211 | 3.58 | 26.32** | 1 | 3 |
| Handicrafts, jewelry, etc | 6 | 0.48 | 78 | 0.71 | 6 | 1.62 | 109 | 1.85 | $3.64^{NS}$ | 0 | 0 |
| Sport hunting/specimen collecting | 26 | 2.07 | 453 | 4.11 | 6 | 1.68 | 168 | 2.85 | $0.12^{NS}$ | 0 | 0 |
| Wearing apparel, accessories | 0 | 0 | 2 | 0.02 | 6 | 1.62 | 217 | 3.69 | $16.21^{NS}$ | 1 | 3 |
| Research | 0 | 0 | 1 | 0.009 | 2 | 0.54 | 35 | 0.59 | $3.10^{NS}$ | 1 | 3 |
| Establishing ex-situ production | 0 | 0 | 1 | 0.009 | 1 | 0.27 | 35 | 0.59 | $0.42^{NS}$ | 2 | 5 |
| Other (free text) | 0 | 0 | 9 | 0.08 | 2 | 0.54 | 26 | 0.44 | $3.10^{NS}$ | 0 | 0 |

**Table S18 Summary result of phylogenetic signal tests which employed to test the threat and population decline of bird and mammal pollinators. The level current indicates the current status of threat or trends of decline. The lower and upper level indicate the treatment of data deficient/unknown population trend species as not threatened/declining or threatened/declining**

| | Bird | | | | | | Mammal | | | | | |
|---|---|---|---|---|---|---|---|---|---|---|---|---|
| | Threat status | | | Population trend | | | Threat status | | | Population trend | | |
| | Current | Lower | Upper | Current | Lower | Upper | Current | Lower | Upper | Current | Lower | Upper |
| Evolutionary Transitions Observed | 84.00 | 84.00 | 86.00 | 322.00 | 368.00 | 359.00 | 52.00 | 54.00 | 63.00 | 61.00 | 74.00 | 90.00 |
| Evolutionary Transitions Randomization Median | 94.00 | 93.00 | 95.00 | 398.00 | 431.00 | 442.00 | 87.00 | 88.00 | 99.00 | 94.00 | 122.00 | 106.00 |
| P-value | 0.001 | 0.001 | 0.001 | 0.001 | 0.001 | 0.001 | 0.001 | 0.001 | 0.001 | 0.001 | 0.001 | 0.001 |

**Table S19 The IUCN Red List status of Critically Endangered and Data deficient bird pollinators of the world from 1988 to 10/05/2024. The population trend shows the current trends of population of these species. Each Red List category indicates: CR = Critically Endangered, EN = Endangered, VU = Vulnerable, NT = Near Threatened, DD = Data Deficient, and NA = Not Available**

| Species | IUCN Status | Population Trend | 1988 | 1994-1999 | 2000-2010 | 2011-2024 | Last Assessed Date |
|---|---|---|---|---|---|---|---|
| *Anthochaera phrygia* | CR | Decreasing | Threatened | EN | EN | CR (EN until 2011) | 06/08/2018 |
| *Ara ambiguus* | CR | Decreasing | NA | EN | EN | CR (EN until 2016) | 07/08/2020 |
| *Campylopterus phainopeplus* | CR | Decreasing | NT | NA | EN | CR | 06/08/2020 |
| *Charmosyna amabilis* | CR | Decreasing | NA | VU | EN/CR | CR | 01/10/2016 |
| *Charmosyna diadema* | CR | Unknown | Threatened | EN | CR | CR | 12/08/2019 |
| *Charmosyna toxopei* | CR | Decreasing | Threatened | VU | CR | CR | 01/10/2016 |
| *Eriocnemis godini* | CR | Unknown | Threatened | CR | CR | CR | 18/08/2020 |
| *Eriocnemis isabellae* | CR | Decreasing | Not recognised | Not recognised | CR | CR | 07/08/2018 |
| *Eulidia yarrellii* | CR | Decreasing | Threatened | VU | EN | CR (EN until 2014) | 11/08/2020 |
| *Gymnomyza aubryana* | CR | Decreasing | NA | VU | EN | CR | 06/08/2018 |
| *Lathamus discolor* | CR | Decreasing | NA | VU | EN | CR | 07/08/2018 |
| *Lophornis brachylophus* | CR | Decreasing | NA | EN | CR | CR | 07/08/2018 |
| *Loxioides bailleui* | CR | Decreasing | Threatened | EN | EN | CR | 20/07/2023 |
| *Oxypogon cyanolaemus* | CR | Decreasing | NA | NA | NA | CR (NA until 2014) | 18/06/2023 |
| *Palmeria dolei* | CR | Decreasing | Threatened | VU | CR | CR | 08/08/2020 |
| *Psittirostra psittacea* | CR | Unknown | Threatened | CR | CR | CR | 01/10/2016 |
| *Sephanoides fernandensis* | CR | Decreasing | Threatened | CR | CR | CR | 15/07/2020 |
| *Vini ultramarina* | CR | Stable | Threatened | EN | EN | CR (EN until 2017) | 09/08/2018 |
| *Zosterops albogularis* | CR | Decreasing | Threatened | CR | CR | CR | 07/08/2018 |
| *Zosterops chloronothos* | CR | Decreasing | Threatened | CR | CR | CR | 01/10/2016 |
| *Zosterops nehrkorni* | CR | Decreasing | Not recognised | Not recognised | CR | CR | 22/08/2018 |
| *Zosterops rotensis* | CR | Increasing | Not recognised | CR | CR | CR | 01/10/2017 |
| *Myzomela albigula* | DD | Unknown | NT | DD | DD | DD | 27/08/2020 |
| *Discosura letitiae* | DD | Unknown | Threatened | DD | DD | DD | 20/09/2022 |

**Table S20 The IUCN Red List status of Critically Endangered and Data deficient mammal pollinators of the world from 1986 to 10/05/2024. The population trend shows the current trends of population of these species. Each Red List category indicates: CR = Critically Endangered, EN = Endangered, VU = Vulnerable, NT = Near Threatened, LR = Lower Risk, LC = Least Concerns, DD = Data Deficient, EX = Extinct and NA = Not Available**

| Species | IUCN Status | Population Trend | 1986-1990 | 1991-1999 | 2000-2010 | 2011-2024 | Last Assessed Date |
|---|---|---|---|---|---|---|---|
| *Brachyteles arachnoides* | CR | Decreasing | EN (EN since 1982) | EN | CR/EN (EN since 2003-2008) | CR | 18/03/2019 |
| *Brachyteles hypoxanthus* | CR | Decreasing | EN | EN | CR | CR | 18/03/2019 |
| *Burramys parvus* | CR | Decreasing | NA | NA/EN (EN since 1994) | EN/CR (CR since 2007) | CR | 30/06/2008 |
| *Callithrix flaviceps* | CR | Decreasing | EN (since 1982) | EN | EN | CR (EN until 2019) | 15/07/2019 |
| *Eulemur mongoz* | CR | Decreasing | VU/EN (EN since 1990) | EN/VU (VU since 1996) | VU | CR (VU until 2013) | 08/05/2018 |
| *Gymnobelideus leadbeateri* | CR | Decreasing | VU (EN since 1982-1986) | EN | EN | CR (EN until 2013) | 22/04/2014 |
| *Mystacina robusta* | CR | Unknown | EX | EX | EX/CR (CR since 2008) | CR | 15/10/2020 |
| *Phyllonycteris aphylla* | CR | Decreasing | NA | EN (EN since 1996) | EN/LC (LC since 2008) | CR (LC until 2014) | 06/03/2015 |
| *Pteropus aruensis* | CR | Unknown | NA | NA | NA/CR (CR since 2008) | CR | 18/01/2016 |
| *Pteropus howensis* | CR | Decreasing | NA | VU | VU/DD (DD since 2008) | CR (DD until 2018) | 28/07/2019 |
| *Pteropus livingstonii* | CR | Decreasing | EN | EN/CR (CR since 1996) | CR/EN (EN since 2008) | CR (EN until 2015) | 31/05/2016 |
| *Varecia variegata* | CR | Decreasing | NA/EN (EN since 1990) | EN/CR (CR since 2008) | CR | CR | 30/12/2019 |
| *Anoura fistulata* | DD | Unknown | NA | NA | NA/DD (DD since 2008) | DD | 03/10/2017 |
| *Artibeus inopinatus* | DD | Unknown | NA | NA/LR/NT (LR/NT since 1996) | VU/DD (DD since 2008) | DD | 29/06/2016 |
| *Epomophorus grandis* | DD | Unknown | NA | Rare/EN (EN since 1996) | DD (DD since 2000) | DD | 20/04/2015 |
| *Lonchophylla chocoana* | DD | Unknown | NA | NA | NA/DD (DD since 2008) | DD | 29/06/2016 |
| *Lonchophylla orcesi* | DD | Unknown | NA | NA | NA/DD (DD since 2008) | DD | 20/07/2015 |
| *Micropteropus intermedius* | DD | Unknown | NA | Rare/VU (VU since 1996) | DD | DD | 23/06/2020 |
| *Platyrrhinus umbratus* | DD | Unknown | NA | NA | NA | DD (DD since 2016) | 28/06/2016 |
| *Pteropus intermedius* | DD | Unknown | NA | NA | NA/DD (DD since 2008) | DD | 25/10/2018 |
| *Pteropus keyensis* | DD | Unknown | NA | NA | NA/DD (DD since 2008) | DD | 18/01/2016 |
| *Pteropus lombocensis* | DD | Unknown | NA | LR (LR since 1996) | LR/DD (DD since 2008) | DD | 18/01/2016 |
| *Pteropus speciosus* | DD | Unknown | Rare (at 1990) | Rare/VU (VU since 1996) | VU/DD (DD since 2008) | DD | 20/10/2018 |
| *Rousettus linduensis* | DD | Unknown | NA | NA | NA/DD (DD since 2008) | DD | 26/05/2019 |
| *Scleronycteris ega* | DD | Decreasing | NA | NA/VU (VU since 1996) | VU/DD (DD since 2008) | DD | 05/08/2016 |
| *Sturnira mistratensis* | DD | Unknown | NA | NA | NA/DD (DD since 2008) | DD | 20/07/2015 |
| *Vampyressa pusilla* | DD | Unknown | NA | NA/LR (LR since 1996) | LR/DD (DD since 2008) | DD | 24/06/2016 |
| *Xeronycteris vieirai* | DD | Unknown | NA | NA | NA/DD (DD since 2008) | DD | 20/07/2015 |

**Table S21 The IUCN Red List status of Critically Endangered, Endangered and Data deficient reptile pollinators of the world from 1986 to 10/05/2024. The population trend shows the current trends of population of these species. Each Red List category indicates: CR = Critically Endangered, EN = Endangered, VU = Vulnerable, DD = Data Deficient and NA = Not Available**

| Species | IUCN Status | Population Trend | 1986-1990 | 1991-1999 | 2000-2010 | 2011-2024 | Last Assessed Date |
|---|---|---|---|---|---|---|---|
| *Cyclura carinata* | EN | Stable | Rare | Rare/CR (CR since 1996) | CR | CR/EN (EN since 2019) | 16/12/2019 |
| *Phelsuma borbonica* | EN | Decreasing | NA | NA | NA | EN (EN since 2019) | 18/12/2019 |
| *Podarcis lilfordi* | EN | Decreasing | VU | VU | VU/EN (EN since 2006) | EN | 14/12/2008 |
| *Toropuku stephensi* | EN | Unknown | VU | VU | VU | VU/EN (EN since 2018) | 17/02/2018 |
| *Hemidactylus dracaenacolus* | CR | Decreasing | NA | NA | NA | CR | 12/07/2011 |
| *Pseudocordylus subviridis* | DD | Unknown | NA | NA | NA | NA | NA |

**Table S22 Patterns of population trends of threatened and not-threatened bird, mammal and reptile pollinators of the world. The table shows the proportion of species that are currently decreasing, stable and increasing along with their lower and upper levels of population trend by considering the Unknown trend species as not-threatened and threatened. N indicates the total number of species of each threatened and not-threatened category**

| **Group** | | *N* | **Unknown** | **Decreasing** | Current | Lower | Upper | **Stable** | Current | Lower | Upper | **Increasing** | Current | Lower | Upper |
|---|---|---|---|---|---|---|---|---|---|---|---|---|---|---|---|
| Birds | Not threatened | 1151 | 145 | 423 | 42.05 | 36.75 | 49.35 | 515 | 51.19 | 44.74 | 57.34 | 68 | 6.76 | 5.91 | 18.51 |
| | Threatened | 104 | 4 | 84 | 84.00 | 80.77 | 84.62 | 13 | 13.00 | 12.50 | 16.35 | 3 | 3.00 | 2.88 | 6.73 |
| Mammals | Not threatened | 269 | 86 | 68 | 37.16 | 25.28 | 57.25 | 110 | 60.11 | 40.89 | 72.86 | 5 | 2.73 | 1.86 | 33.83 |
| | Threatened | 102 | 9 | 84 | 90.32 | 82.35 | 91.18 | 7 | 7.53 | 6.86 | 15.69 | 2 | 2.15 | 1.96 | 10.78 |
| Reptiles | Not threatened | 35 | 10 | 1 | 4.00 | 2.86 | 31.43 | 23 | 92.00 | 65.71 | 94.29 | 1 | 4.00 | 2.86 | 31.43 |
| | Threatened | 5 | 1 | 3 | 75.00 | 60.00 | 80.00 | 1 | 25.00 | 20.00 | 40.00 | 0 | 0.00 | 0.00 | 20.00 |

**Table S23 Patterns of population trends of threatened endemic bird, mammal and reptile pollinators of the world. The table shows the proportion of species that are currently decreasing, stable and increasing along with their lower and upper levels of population trend by considering the Unknown trend species as not-threatened and threatened. N indicates the total number of species of each group**

| | IUCN Population Trends | | | | | | | | | | | | | |
|---|---|---|---|---|---|---|---|---|---|---|---|---|---|---|
| **Group** | *N* | **Unknown** | **Decreasing** | Current | Lower | Upper | **Stable** | Current | Lower | Upper | **Increasing** | Current | Lower | Upper |
| Birds | 93 | 4 | 73 | 82.02 | 78.49 | 82.80 | 13 | 14.61 | 13.98 | 18.28 | 3 | 3.37 | 3.23 | 7.53 |
| Mammals | 56 | 3 | 48 | 90.57 | 85.71 | 91.07 | 3 | 5.66 | 5.36 | 10.71 | 2 | 3.77 | 3.57 | 8.93 |
| Reptiles | 4 | 1 | 3 | 100.0 | 75.00 | 100.0 | 0 | 0.00 | 0.00 | 25.00 | 0 | 0.00 | 0.00 | 25.00 |

*Figures*

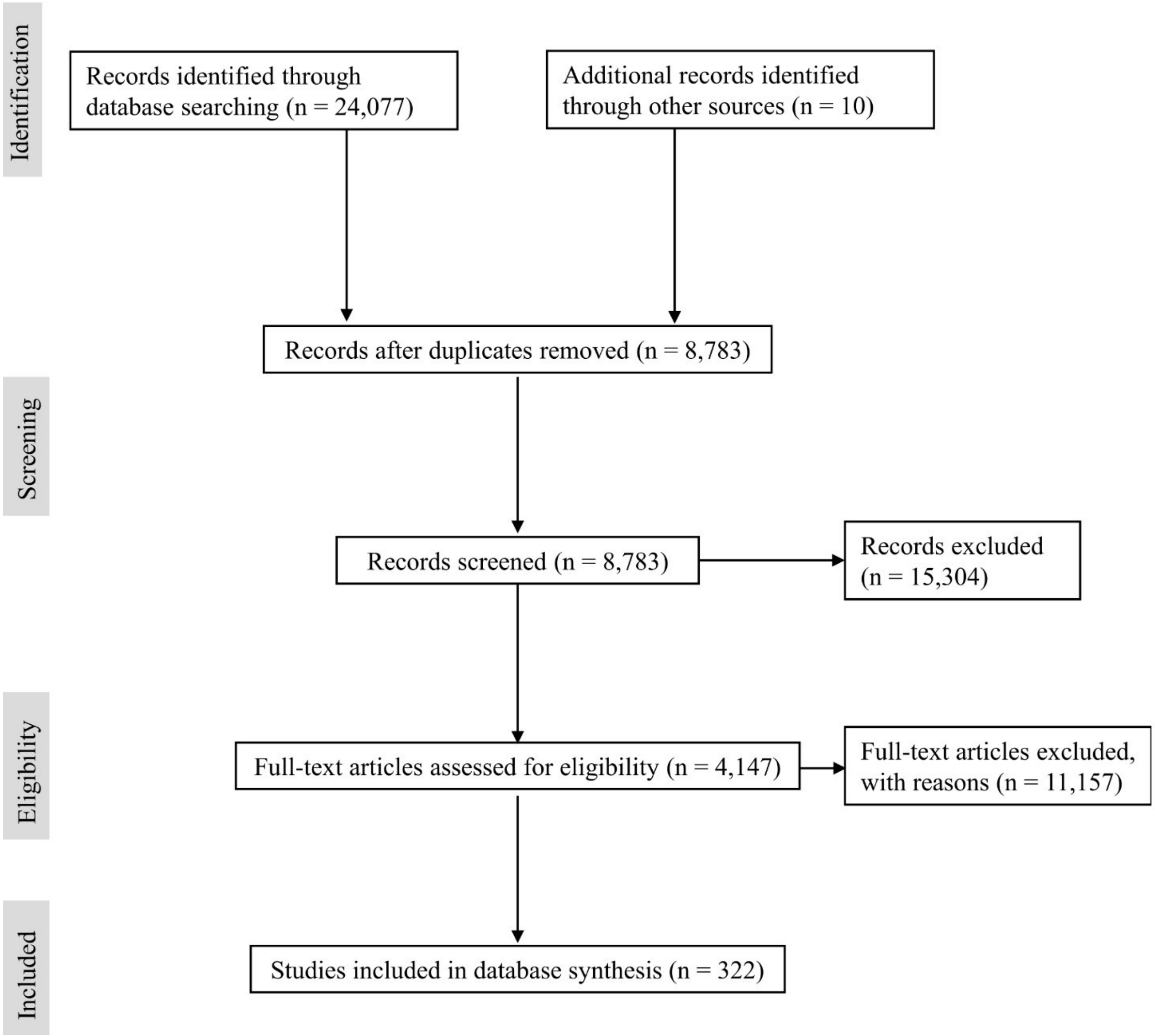


**Figure S1 Preferred Reporting Items for Systematic Review and Meta-Analysis flowchart (PRISMA), summarizing the sequence of information gathering and selection of vertebrate pollinator studies.**

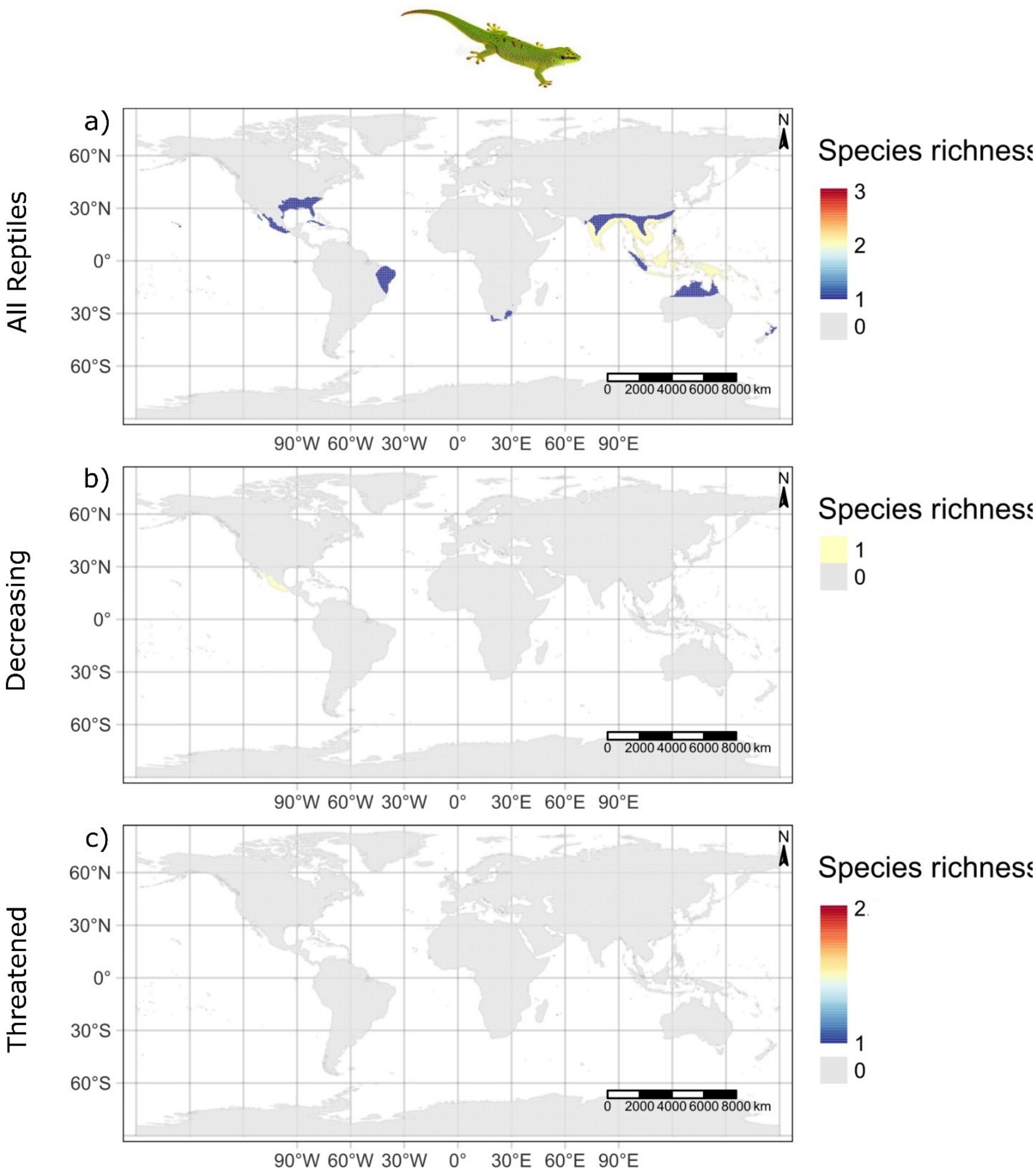


**Figure S2 The spatial richness patterns of (a) all (b) decreasing and (c) threatened reptile pollinators of the world. A spatial resolution of 0.5-degree latitudinal and longitudinal grids (approximately 50 km) was used to create maps**

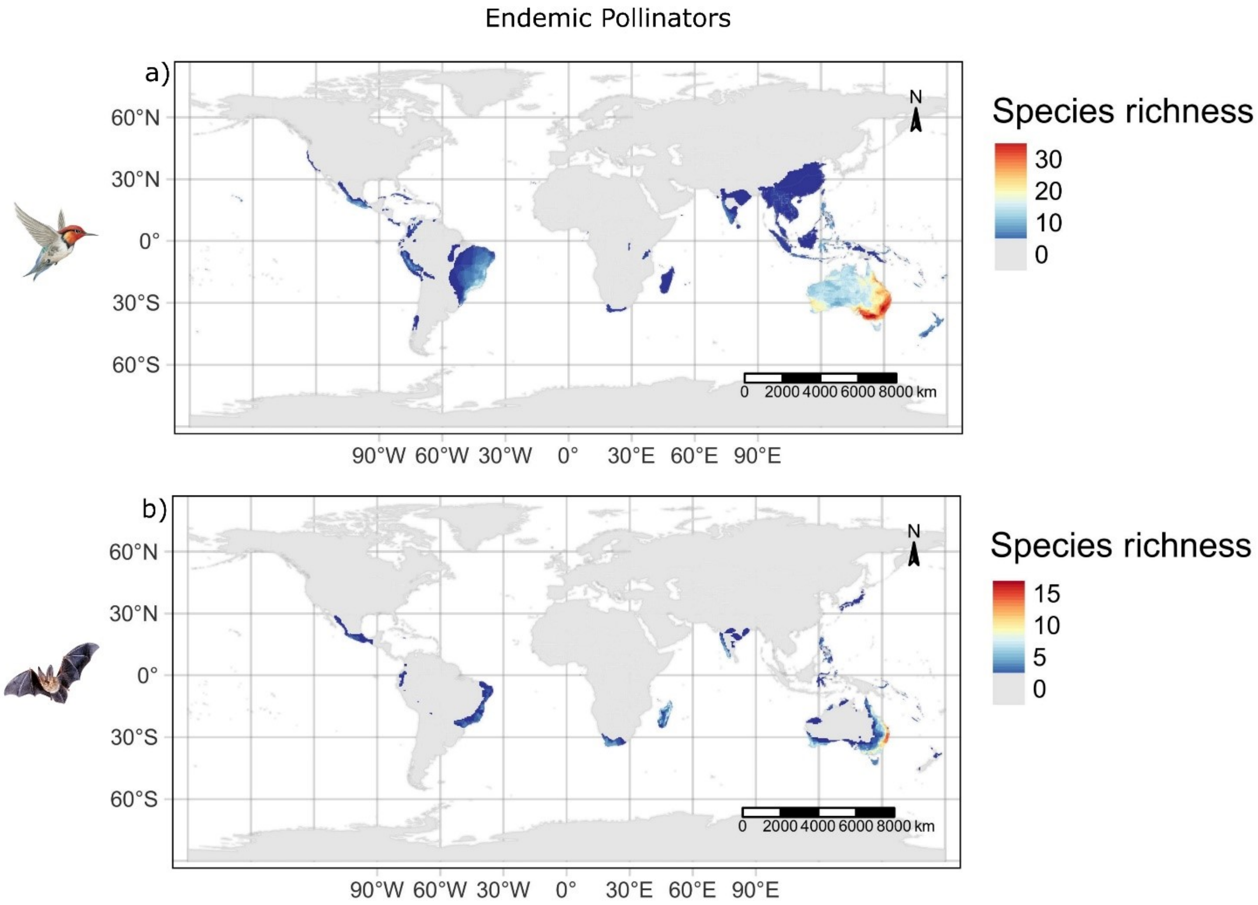


**Figure S3** **The spatial richness pattern of endemic (a) bird and (b) mammal pollinators of the world. A spatial resolution of 0.5-degree latitudinal and longitudinal grids (approximately 50 km) was used to create maps**

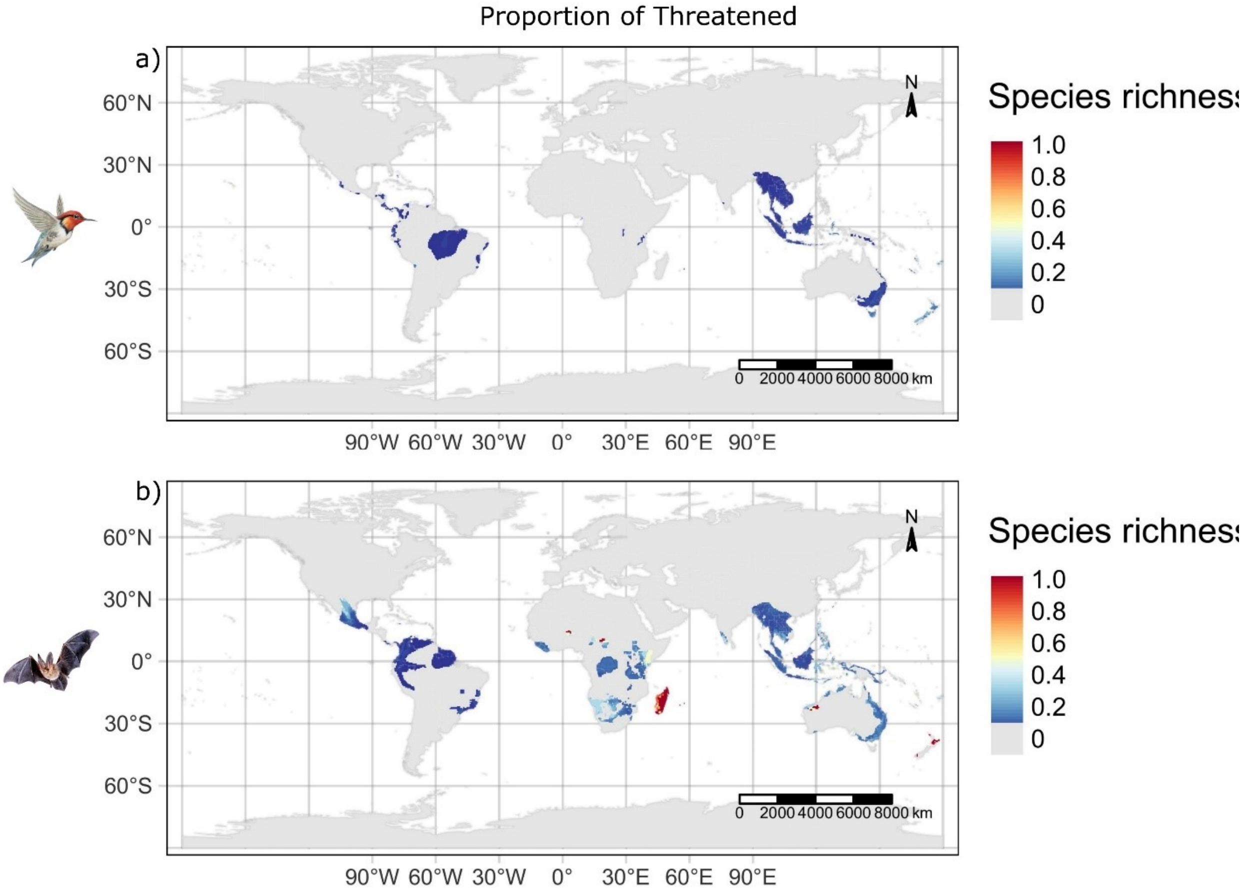


**Figure S4 The spatial richness patterns of proportion of threatened (a) bird and (b) mammal pollinators of the world. A spatial resolution of 0.5-degree latitudinal and longitudinal grids (approximately 50 km) was used to create maps**

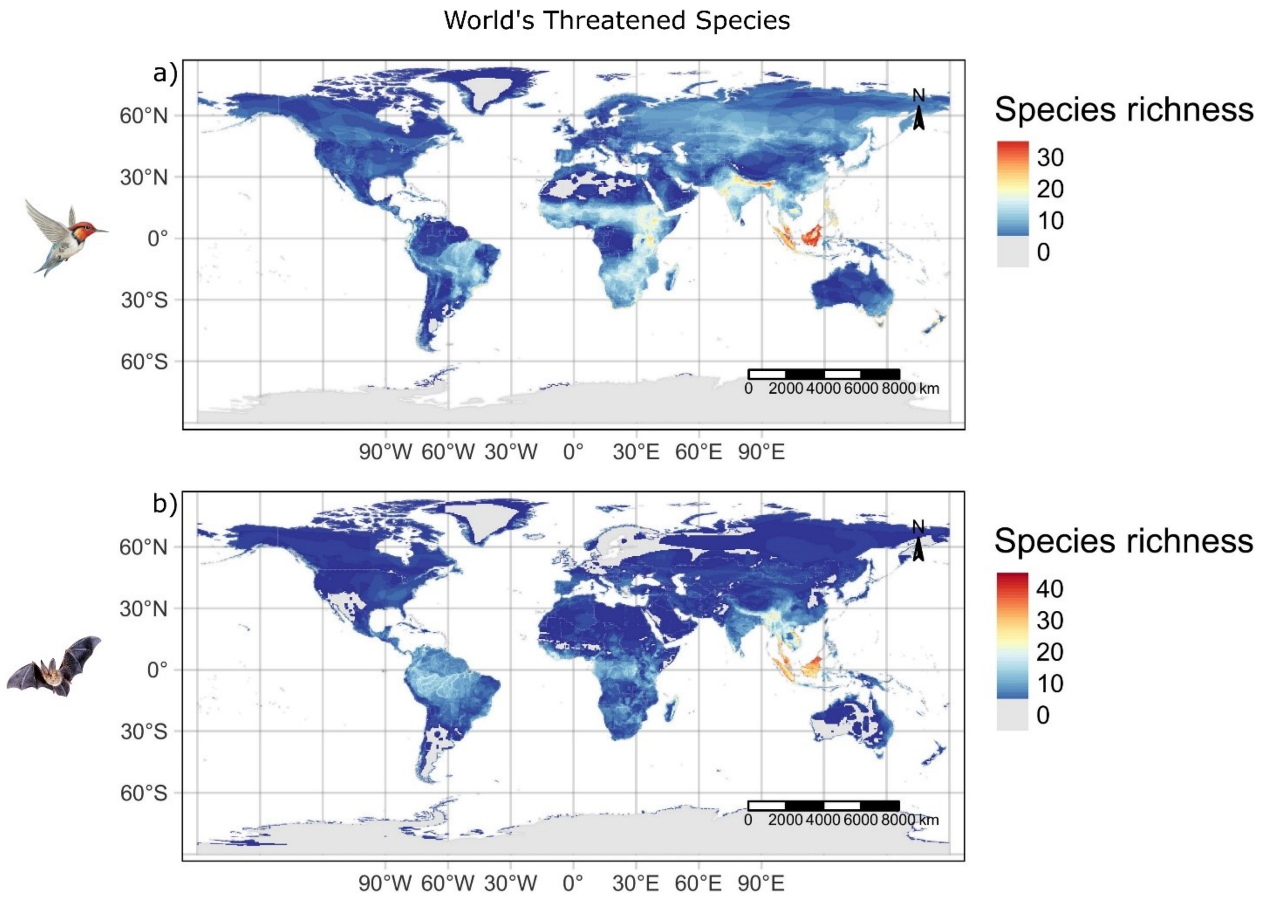


**Figure S5 The spatial richness patterns of generalist (a) bird and (b) mammal species of the world. A spatial resolution of 0.5-degree latitudinal and longitudinal grids (approximately 50 km) was used to create maps**

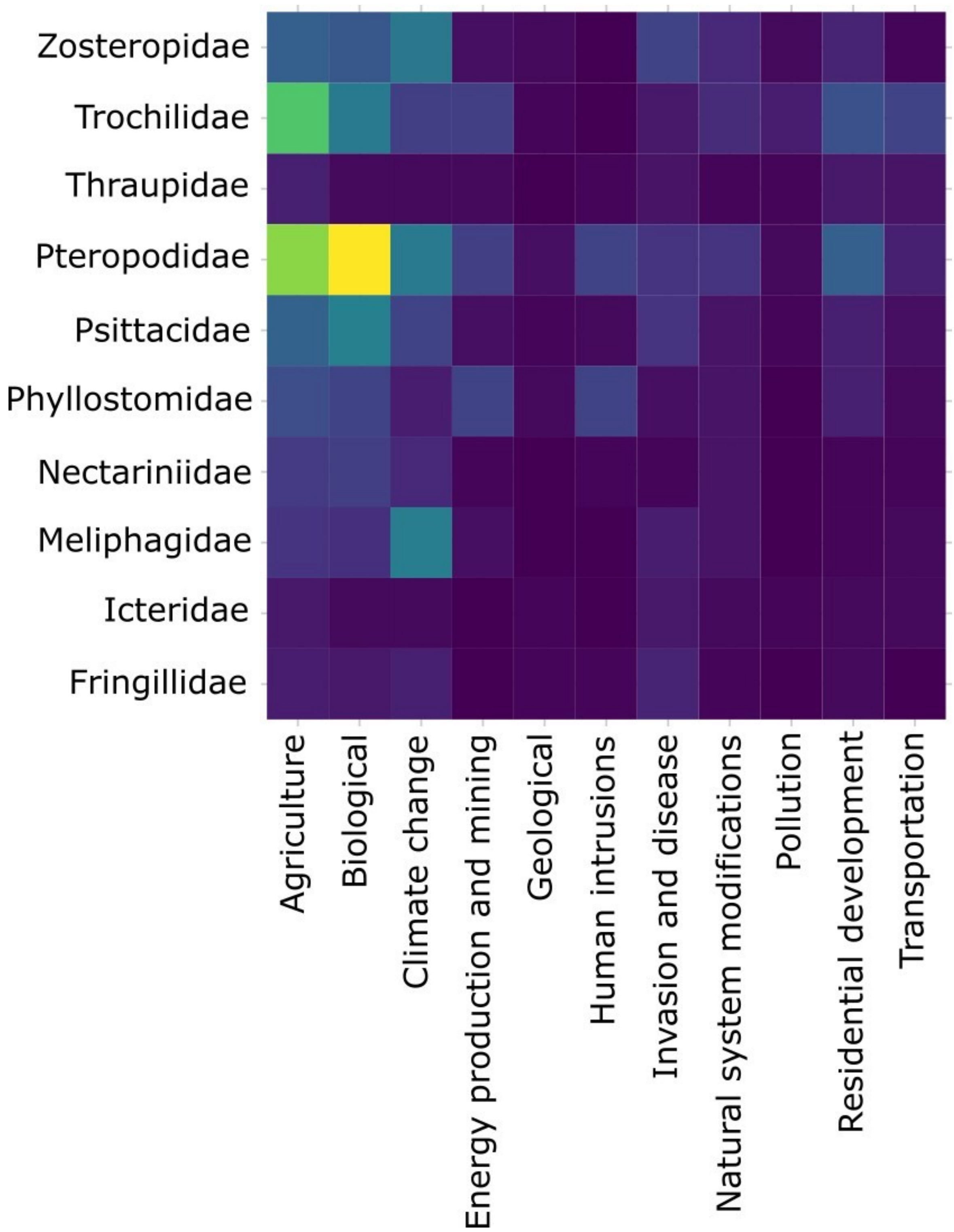


**Figure S6** **Various types of anthropogenic threats on major families of bird and mammal pollinators. The categorization of major family for birds was based on the family that contains minimum of 27 or more species. For mammals, this was based on the family that contains a minimum of 100 species**

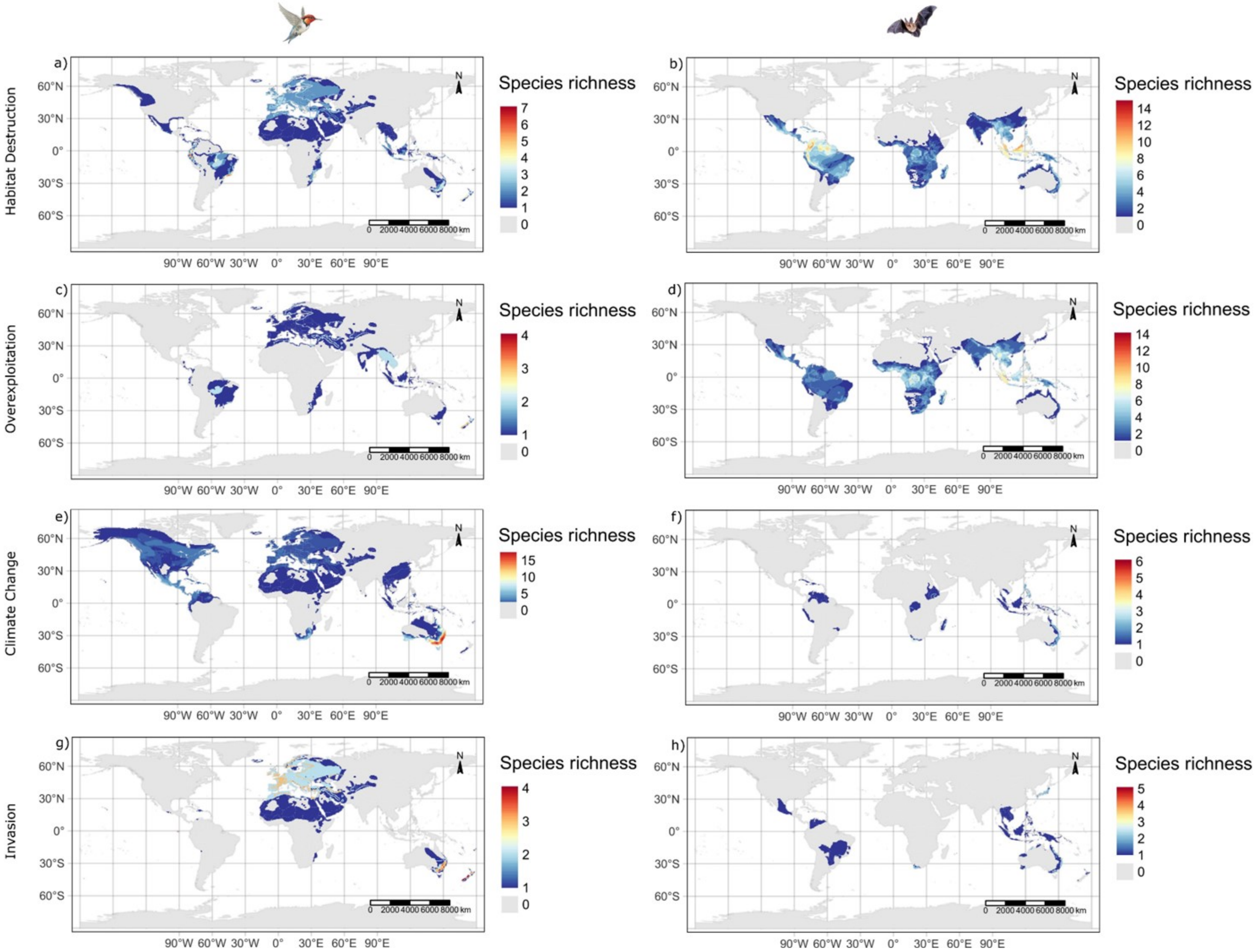

a)
b)
c)
d)
e)
f)
g)
h)
Habitat Destruction
Overexploitation
Climate Change
Invasion
Species richness
7
6
5
4
3
2
1
0
14
12
10
8
6
4
2
0
4
3
2
1
0
15
10
5
0
6
5
4
3
2
1
0
4
3
2
1
0
5
4
3
2
1
0
60°N
30°N
0°
30°S
60°S
90°W 60°W 30°W 0° 30°E 60°E 90°E
N
0 2000 4000 6000 8000 km

**Figure S7 The spatial richness patterns of (a, c, e, g) bird and (b, d, f, h) mammal pollinators that are experiencing various threats from human activities (anthropogenic factors). A spatial resolution of 0.5-degree latitudinal and longitudinal grids (approximately 50 km) was used to create maps**

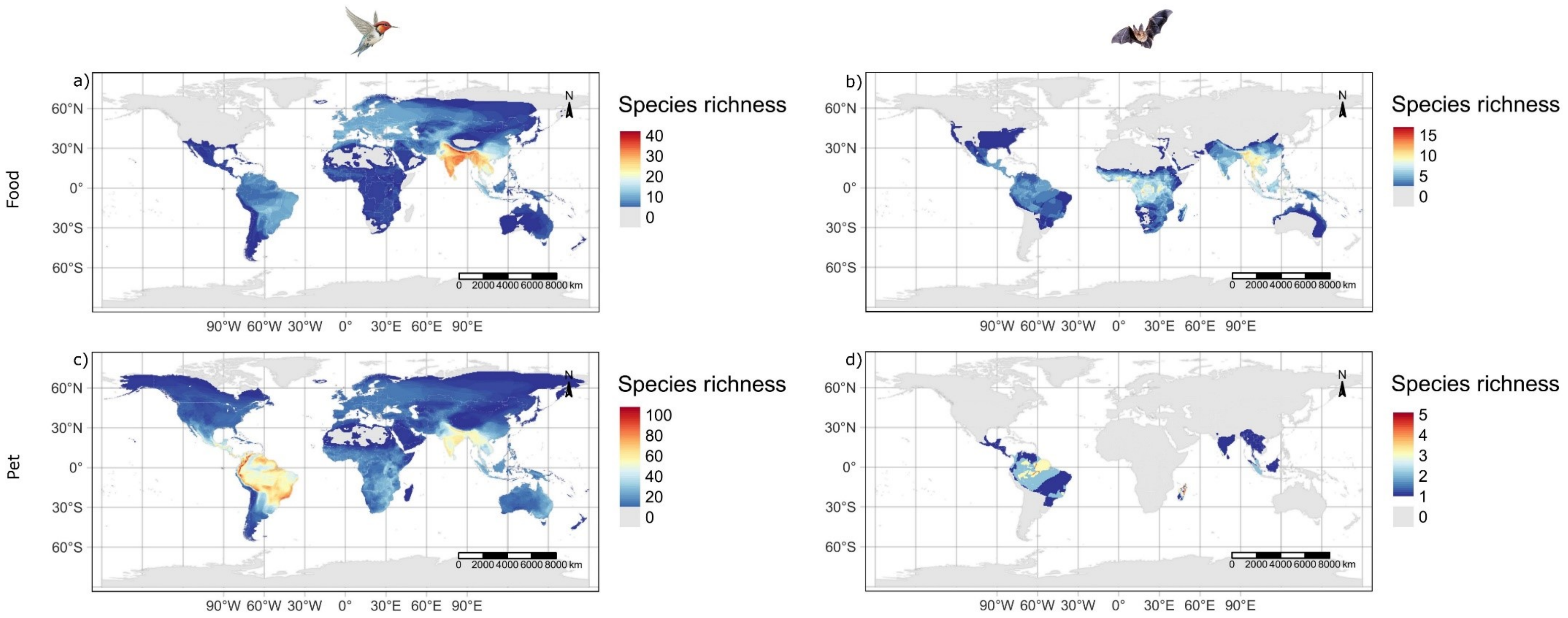


**Figure S8 The spatial richness patterns of (a, c) bird and (b, d) mammal pollinators that used in the (a, b) food and (c, d) pet markets across the world. A spatial resolution of 0.5-degree latitudinal and longitudinal grids (approximately 50 km) was used to create maps**

#### *Reference*


Ali, S. (1931). The role of sunbirds and flowerpeckers in the propagation and distribution of the tree-parasite, Loranthus longiflorus Dest., in the Konkan (W. India). J. Bombay nat. Hist. Soc., 35, 144-149

Davidar, P. (1985). Ecological interactions between mistletoes and their avian pollinators in south India. Journal of the Bombay Natural History Society.

Diller, C., Castañeda-Zárate, M. and Johnson, S.D., 2019. Generalist birds outperform specialist sunbirds as pollinators of an African Aloe. Biology letters, 15(7), p.20190349.

Faegri, K. and Van Der Pijl, L., 2013. Principles of pollination ecology.

McCann, C. (1931). On the fertilisation of the flowers of the sausage tree (Kigelia pinnata) by bats. Journal of Bombay Natural History Society, 35, 467-471.

McCann, C. (1933). The flying fox (P. giganteus) and the palm squirrel (F. tristiatus) as agents of pollinization in (Grevillea robusta A. Cunn.) the silky oak. Journal of Bombay Natural History Society, 36, 761-764.

Nicodemus, A., J. Prasanth Jacob, and B. Nagarajan. "Pollination by nectarivorous birds in teak clonal seed orchards." (2006): 1613-1616.

Proctor, M., Yeo, P. and Lack, A., 1996. The natural history of pollination. HarperCollins Publishers.

Raju, A.S., 1990. Observations on the floral biology of certain mangroves. *Proceedings of the Indian National Science Academy. Part B, Biological Sciences*, *56*(4), pp.367-374.

Raju, A.S., 2009. pollination biology and sustainability of economically important acacia sinuata (shikai) and semecarpus anacardium (marking nut) in the eastern ghats forests of Andhra Pradesh, India. *Bioresources Conservation and Management*, *17*, p.199.

Singh, T. (1929). A preliminary note on the pollination of the coral tree (Erythrina indica Lamk.). Bombay Nat. Hist. Soc, 33, 460-462.
